# Structural mapping of patient-associated *KCNMA1* gene variants

**DOI:** 10.1101/2023.07.27.550850

**Authors:** Hans J. Moldenhauer, Kelly Tammen, Andrea L. Meredith

**Affiliations:** Dept. of Physiology, University of Maryland School of Medicine, Baltimore, MD, USA

**Keywords:** BK channel, KCa1.1, calcium-activated potassium channel, *KCNMA1*, potassium channel, Slo1, slowpoke, channelopathy, pathogenicity algorithm

## Abstract

*KCNMA1*-linked channelopathy is a neurological disorder characterized by seizures, motor abnormalities, and neurodevelopmental disabilities. The disease mechanisms are predicted to result from alterations in *KCNMA1*-encoded BK K^+^ channel activity; however, only a subset of the patient-associated variants have been functionally studied. The localization of these variants within the tertiary structure or evaluation by pathogenicity algorithms has not been systematically assessed. In this study, 82 nonsynonymous patient-associated *KCNMA1* variants were mapped within the BK channel protein. Fifty-three variants localized within cryo-EM resolved structures, including 21 classified as either gain-of-function (GOF) or loss-of-function (LOF) in BK channel activity. Clusters of LOF variants were identified in the pore, the AC region (RCK1), and near the Ca^2+^ bowl (RCK2), overlapping with sites of pharmacological or endogenous modulation. However, no clustering was found for GOF variants. To further understand variants of uncertain significance (VUS), assessments by multiple standard pathogenicity algorithms were compared, and new thresholds for sensitivity and specificity were established from confirmed GOF and LOF variants. An ensemble algorithm was constructed (KCNMA1 Meta Score), consisting of a weighted summation of this trained dataset combined with a structural component derived from the Ca^2+^ bound and unbound BK channels. KMS assessment differed from the highest performing individual algorithm (REVEL) at 10 VUS residues, and a subset were studied further by electrophysiology in HEK293 cells. M578T, E656A, and D965V (KMS^+^;REVEL^-^) were confirmed to alter BK channel properties in voltage-clamp recordings, and D800Y (KMS^-^;REVEL^+^) was assessed as benign under the test conditions. However, KMS failed to accurately assess K457E. These combined results reveal the distribution of potentially disease-causing *KCNMA1* variants within BK channel functional domains and pathogenicity evaluation for VUS, suggesting strategies for improving channel-level predictions in future studies by building on ensemble algorithms such as KMS.

## Introduction

*KCNMA1*-linked channelopathy is a rare, predominantly neurological syndrome of incompletely defined phenotypes associated with mutations in the *KCNMA1* gene (1). The protein product of the *KCNMA1* gene is the pore-forming subunit of the calcium and voltage-activated BK K^+^ channel (2-4). BK channels regulate neuronal, muscle, and neuroendocrine cell excitability, with additional roles in non-excitable cells in tissues such as kidney, bone, and gastrointestinal tract (2,5-7). Overlapping symptomology between patients has identified four main dysfunctions in individuals with *KCNMA1* variants: epilepsy, movement disorder, neurodevelopmental delay and intellectual disability, and structural malformations (8,9). At present, two variants (D434G and N999S) are conclusively determined to cause neurological disease, based on co-segregation of seizure and paroxysmal nonkinesigenic dyskinesia (PNKD) in a family pedigree and recapitulation of those phenotypes in animal models (10-13). However, there is high phenotypic variability across variants, and the majority of human patient (HP)-associated variants are de novo and found only in a single individual. Moreover, it is currently unclear whether all *KCNMA1* variants discovered through genetic testing are causative. Both functional and computational studies assessing the pathogenicity are incomplete.

The goal of this study was to systematically determine the location and report the pathogenicity associated with *KCNMA1* HP variants. The study was restricted to HP variants and did not include other types of engineered mutations known to alter gating, which prioritizes mutations with disease-causing potential. In the first step, we undertook a comprehensive annotation of the cryo-EM structure of the tetrameric BK channel pore-forming subunit to probe the domain-based locations for HP variants and assessed whether there was evidence for clustering of pathogenic variants. In the second step, we assessed the performance of pathogenicity algorithms on HP variants, starting with standard ClinVar algorithms which lack BK channel-specific validity. We next trained a subset of these algorithms on *KCNMA1* HP variants of known pathogenicity confirmed by functional data and integrated structural parameters from open and closed BK channel conformations into an ensemble algorithm (*KCNMA1* Meta Score). KMS analysis identified several VUS where the pathogenicity assessment differed from REVEL, an algorithm without BK channel-specific training. These variants were assessed for functional effects on BK channel properties using patch-clamp electrophysiology in heterologous cells.

## Methods

### Linear Sequence Annotation

*KCNMA1* variants, named according to the original publications or clinical reports, were annotated in the human BK channel sequence MG279689 (Supplemental Figure 1). This wildtype (WT) isoform has the alternative exons SRKR (site 1), no insert (site 2; STREX), Ca^2+^ bowl (site 3), and VYR (site 4; C-term). Variants were obtained from ClinVar, the Sanford Coordination of Rare Diseases (CoRDs) Registry, IRB protocol #03-10-014 (Sanford Research, University of South Dakota), and direct submissions to ALM under University of Maryland School of Medicine IRB Non-Human Subject Research (NHSR) designations HP-00083221 and HP-00086440. Single nucleotide polymorphisms (SNPs) were obtained from gnomAD.

### Structural and cluster analysis

Distance between residues was calculated in Visual Molecular Dynamics (14), measuring between alpha carbons. A cluster was defined as three residues or more residues in a domain within 5Å of each other. This distance is considered the average interaction distance in energetic terms (15). For reference, the gating ring measures 43.5 Å across in the open state between K384 α carbons of diagonal subunits, and the channel’s length is 105.6 Å between Q19 in the N terminal and M712 in the RCK2. Residue pairs were defined with these same parameters, but only between two residues. The analysis was performed on the cryo-EM BK channel Ca^2+^ open channel structure (PDB ID:6V38) and cryo-EM BK channel Ca^2+^ free closed channel structure (PDB ID: 6V3G) (16). Since no change in classifications of pairs and clusters were observed, the calcium bound channel was utilized for representation in each figure. This cryo-EM structure spans amino acid 19 to 1056 and lacks the regions between the amino acids: 51 to 93, 613 to 684, and 833 to 871 (Supplemental Figure 1). The VMD visualization settings were ‘cartoon’ for the complete channel structure and ‘Van der Waals (VDW)’ for representation of individual residues of interest. Additionally, four patient variants corresponding to channel truncations were generated within the cryo-EM structure (PDB ID:6V38) (16) and compared with two experimentally truncated channels: Slo1C truncates in the linker between S6 and RCK1 at position 342 and Slo1ΔRCK2 truncates after RCK1 at position 632 (17). The Slo1C representation does not include the 74 amino acid tail, or the 11 amino acid residues from Kv1.4, that were included in the published Slo1C construct. The Slo1ΔRCK2 structural model was used as reference for the RCK1 and RCK2 boundaries for comparison to other representation truncation variants. Nonsynonymous single nucleotide polymorphisms (SNPs) from GenomAD (ENSG00000156113.16) (8, 25) were added onto the cryo-EM structure (PDB ID: 6V38).

### Pharmacological and Intracellular Modulators

Thirteen of sixteen modulators (agonist, antagonist, and endogenous molecules) with action on the alpha subunit of the BK channel were mapped. The agonists included were BC5 (18), NS1619 (52, 53) and Cym04 (52). The antagonists were charybdotoxin (ChTX) (46), iberiotoxin (IbTX) (47), loperamide (LOP) (48), paxilline (PAX) (49), and TEA (50, 51). The endogenous modulators were Heme (54), protons (H^+^) (55), lipids (56), carbon monoxide (CO) (57), kinases (58) and ethanol (EtOH) (59). Phospho-sites are numbered in the context of the BK channel cDNA NM_001161352.2. Only modulators with confirmed or putative binding sites were included.

### Pathogenicity scores

The algorithms SIFT, Polyphen, CADD/PHRED, MetaLR, M-CAP, Mutpred and REVEL were evaluated in their predictor value for the *KCNMA1* patient mutations using the online variant predictor tool from www.ensembl.org. Default settings were used with the *KCNMA1* reference sequence number associated with the human patient variants reported in Clinvar database: NM_001161352.2. Pathogenic designations were based on scores above either standard or experimentally-determined cutoff values. The parameters evaluated for experimentally-determined cutoffs were accuracy (how close the pathogenic scores are from confirmed ones), precision (how close the pathogenic variant scores are), specificity (ability to detect a benign variant correctly), and sensitivity (ability to identify a pathogenic variant correctly) according to the equations 1 to 4 using the pathogenicity scores obtained for the confirmed benign and pathogenic variants (variants listed in Supplemental Table 1B):

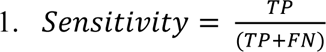

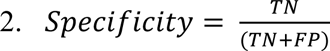

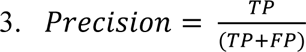

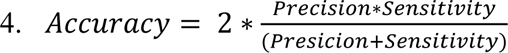

With the abbreviations of TP being true positive, TN being true negative, FP being false positive and FN being false negative (Supplemental Table 1). The numbers associated to each term vary between algorithms and are based on the confirmed pathogenic and non-pathogenic variants. The total performance score for each algorithm was calculated averaging the 4 parameters to determine the goodness series to determine its inclusion in the *KCNMA1* meta score algorithm construction (Figure 6) (19).

For REVEL, CADD/Phred, Mutpred, MetaLR, and WSM, pathogenicity cutoff determinations for significance were made using a training data set with 16 pathogenic variants confirmed by experimental data (A172T, S351Y, G354S, G356S, G375R, C413Y, D434G, H444Q, N536H, G567S, I663V, P805L, D984N, N999S, R1083K, R1097H) assigned value of 1, and 4 benign variants (K518N, E884K, R1128W and N1159S) assigned value of 0. Cutoffs for each algorithm were determined from the training data set using the ‘the closest to (0,1) criteria’ (20). Receiver operating characteristic (ROC) curves, visualizing the sensitivity versus (1 – specificity) for all possible cutoffs were generated in Prism 9 (GraphPad Software, San Diego, CA; Figure 6C insets). The lowest false positive and the highest true positive value was used to determine the algorithm cutoff from the logistic regression fit in pathogenicity score plots (Prism 9; Figure 6C main graphs). Values greater than the cutoff were considered significant.

### *KCNMA1* meta score (KMS) algorithm

To construct the KMS algorithm, a weighted summation model (WSM) that incorporates structural information (19,21) from REVEL, Mutpred, MetaLR and CADD/PHRED was used. REVEL, Mutpred, and MetaLR algorithms scores ranged from 0 to 1, and were not normalized. The CADD/Phred score was divided by 10 to avoid over-representation in the WSM (equation 5; Results section). The structural component (SC) was calculated from the human BK channel cryo-EM structures in absence and presence of Ca^2+^ (PDB:6V3G and 6V38) (16,22). The SC consist in the interactions and the energy associated between residues, calculated using the web based algorithm RING with the default settings (https://ring.biocomputingup.it/submit) (23) to determine the number of interactions (NI; hydrogen bonds, salt bridge, Π-cation, Π-Π stacking, Van der Waals, and disulphide bonds) and average energy (AE). The number of residues (NR) was determined using the ProteinTools Contact Map web tool to calculate the distance between each patient variant residue and all the other residues in the cryo-EM structures. 5Å was used as the cutoff value determined from the number of residues surrounding the residue of interest (https://proteintools.uni-bayreuth.de) (24). NI, AE, and NR values are presented in Supplemental Table 2. Cutoffs for KMS were determined with the same training data set of 16 pathogenic and 4 benign residues.

### Synthesis of *KCNMA1* patient variants in the human BK channel

The variants D800Y, D965V, E656A and M578T were introduced by site-directed mutagenesis into the WT human BK channel in pcDNA3.1+ (Bioinnovatise, Rockville, MD) and verified by sequencing. This BK channel cDNA includes Myc at the N-terminus and the EYFP tag inserted at exon 22, at position 716 in RCK2 (25). YFP fluorescence was used to identify transfected cells. These tags do not affect BK current properties under these conditions (25,26) and have been used in our previous studies (11,25,27-29).

### Electrophysiology

HEK293T cells were used to express the BK channel constructs (CRL-11268, ATCC, Manassas, VA, USA). The cell culture was incubated at 37°C with 5% carbon dioxide. The cells were plated on 35 mm tissue culture dish and fed with DMEM containing media (Cat. #11995-065, Gibco, Life Technologies Corp., Grand Island, NY, USA) supplemented with: 10% fetal bovine serum (Cat. #4135, Sigma-Aldrich, St. Louis, MO, USA), 1% penicillin/streptomycin (Cat. #30-002-Cl, Mediatech Inc., Manassas, VA, USA) and 1% L-glutamine (Cat. #25-005-Cl, Mediatech Inc., Manassas, VA, USA). Cells were transfected at 60-70% confluency using Trans-IT LT1 (Mirius Biological, Madison, WI, USA) at 1:2 ratio of cDNA to transfection reagent (μg/μL). After 6 hours, cells were re-plated onto poly-L-lysine glass coverslips (Cat. #P4832, Sigma-Aldrich, St. Louis, MO, USA). The recordings were performed at 14-24 hours post-transfection.

Electrophysiology solutions approximated the physiological K^+^ gradient with 10 μM intracellular free Ca^2+^. The external (pipette) solution contained (in mM): 134 NaCl, 6 KCl, 1 MgCl_2_, 10 glucose, and 10 HEPES with pH adjusted to 7.4 with NaOH. The internal (bath) solution contained in mM: 110 KMeSO_3_, 10 NaCl, 30 KCl, 10 HEPES, 1 MgCl_2_, 5 HEDTA and 10 µM CaCl_2_. The pH was adjusted to 7.2 with KOH. The desired nominal Ca^2+^ concentration was achieved using WebMaxC to calculate the volume of a 1 M CaCl_2_ stock (Thermo fisher) in HEDTA (http://web.stanford.edu/~cpatton/webmaxcS.htm).

Macroscopic currents were recorded in inside-out patch clamp configuration at room temperature (22°C to 24°C). Electrodes were made from thin-walled borosilicate glass pipettes, with resistances of 2-3.5 MΩ (Cat. #TW150F-4, World Precision Instruments, Sarasota, FL, USA). Voltage commands were applied with a MultiClamp 700B amplifier (Axon Instruments, Sunnyvale, CA, USA) through a Digidata 1440/1550B interface with CLAMPEX 10/11 suite (Molecular devices). Three current traces were acquired at 50 kHz, filtered online at 10 kHz, and averaged offline. Two voltage step protocols were used for activation and deactivation, respectively. The activation protocol stepped from a V_hold_ of −150 mV to +150 mV (Δ+10 mV increments) for 50 ms and back to −150 mV for 15 ms to generate tail currents. Conductance (G) was calculated from tail currents, 150-200 µs after the peak, and normalized to the highest conductance calculated for each patch (G/G_max_). G/G_max_ values were plotted against membrane potential (V) to generate conductance-voltage (G/G_max_-V) curves. The voltage of half maximal activation (V_1/2_) values were calculated by fitting the G/G_max_-V relationship to a Boltzmann function using Prism v9, constrained to 1 at top and 0 at bottom (GraphPad Software, San Diego, California USA). The deactivation time constant (τ_deact_) were obtained from BK currents activated by 20 ms voltage steps of +200 mV from a V_hold_ of -150 mV, followed by 15 ms voltage steps from -200 mV to -50 mV (Δ+10 mV increments). Activation time constants (τ_act_) were obtained from the voltage activation protocol, by fitting a single standard exponential to the rising phase of the current. Deactivation kinetics were measured by fitting the tail current decay with a single exponential function in pCLAMP 10.6 (Molecular Devices, San Jose, CA, USA). In all protocols, leak current was subtracted using a P/5 protocol with a subsweep V_hold_ of -120 mV. Mutated BK channels were categorized as GOF if the currents showed a statistically significant V_1/2_ shift to hyperpolarized potentials, faster activation kinetics, and slower deactivation kinetics compared to WT currents. LOF BK channels had statistically significant changes in the parameters in the opposite direction. Variants with no statistically significant changes remained categorized as VUS.

### Statistics

BK current analysis was performed with pClamp 10.6 (Molecular Devices, San Jose, CA, USA). Graphs and statistical analysis were generated in Prism v9 (GraphPad Software, San Diego, CA, USA). Two-way ANOVA with Bonferroni post-hoc tests were used to determine significance for V_1/2_ and activation/deactivation kinetics at p < 0.05, and the p values are listed in the figure legend.

## Results

### Locations and clustering of LOF and GOF *KCNMA1* variants

The *KCNMA1* gene contains >1000 variants, 791 of which are within exons and 277 within introns (Supplemental Figure 2). This study reports analysis of 82 nonsynonymous *KCNMA1* human patient (HP)-associated variants from 109 individuals with heterogeneous neurological symptoms (1,8,9)(ALM unpublished data). The names of the variants throughout the text are used from the numbering schemes of the original reports. Fifty-three variants occurred within resolved regions of the human Ca^2+^ bound BK channel cryo-EM structure (Figure1 and Supplemental Figure 1). Nine variants were located within unresolved regions in the cryo-EM structure or occurred in alternate exons and were not analyzed. Thirteen of these variants produced deletions or frameshifts. To understand the structural relationships for variants altering single amino acids with the BK channel, the 53 variants were assessed with respect to the Ca^2+^ binding sites, Mg^2+^ binding site, voltage sensor domain (VSD), selectivity filter residues, pharmacological modulator binding sites, and post-translational modifications (Figure 1). Of these, 15 were categorized as loss of function (LOF), 3 as gain of function (GOF), and 4 as benign from functional studies of BK currents and/or expression. The remaining variants of uncertain significance (VUS) that have not yet been studied functionally were either obtained from individuals with neurological disease or ClinVar (Supplemental Figure 2).

**Figure 1.**
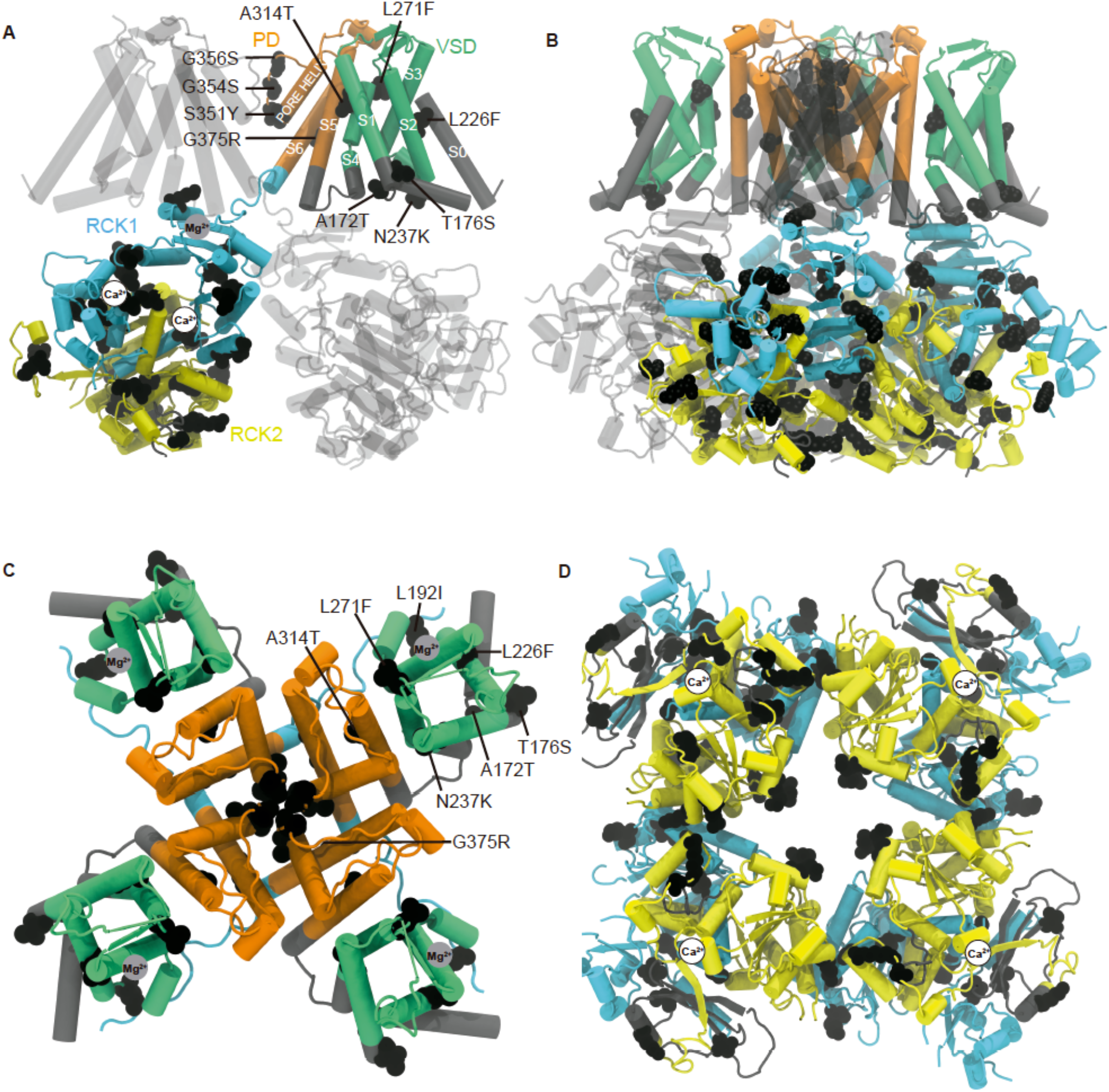
Structural localization of *KCNMA1* HP variants. (A) Fifty-three patient-associated *KCNMA1* variants (black globular residues) are depicted in the calcium-bound human BK channel alpha subunit, with either two (A) or four subunits (B); Cryo-EM PDB ID: 6V38). Voltage sensing domain (VSD; green), pore domain (PD; orange), gating ring (RCK1: light blue and RCK2: yellow), and Ca^2+^ (white circles) and Mg^2+^ binding sites (gray circle). (C-D) Top (C) and bottom (D) view of the BK channel tetramer. Residues 1-18, 51-93, 614-683, 834-870 and 1057-1111 are not shown due to location in unresolved or deleted segments. Residues in the transmembrane domains are labelled in panels A and C.

Missense HP variants were found in exons comprising the three main channel modules: pore domain (PD), voltage sensing domain (VSD) and the gating ring (Figure 1A and B). Interestingly, only 10 HP variants were in the transmembrane segments (Figure 1C top view), corresponding to 2.5% of the residues in this region, compared to 43 located in the intracellular part of the channel (the intracellular linkers and gating ring, 5% of the total residues) (Figure 1D bottom view). HP variants A172T, T176S, L226F, N237K, L271F and A314Y located in the VSD are at distances above 5Å from the residues involved in the voltage activation R178, D198, D218, R232, E245, D251, R272, R275, R278 and E284 (30). The only exceptions were A172 and T176S (3.3 and 3.2Å from R178 in S1) and L271F (4.8Å from R275 in S4).

Next, we determined the clustering of HP variants by localizing the confirmed LOF and GOF HP mutations in the BK channel structure (Figure 2A). A cluster was defined as three residues within 5Å in a domain. In the case of the LOF variants, fifteen of twenty-one variants were placed onto the BK channel cryo-EM structure. Four terminations R458Ter, R640Ter, R830Ter, R880Ter, and the two frameshift N449fs and Y676fs, were excluded since the electrophysiological phenotypes may be due to the sequence truncation or alteration and not to mutation residue itself. In a detailed examination of the LOF variants, two clusters and two pairs were observed. Only three of the fifteen LOF variants, A172T, N237K, and R1097H, did not fall within a pair or cluster (Figure 2A). One of the clusters of LOF variants occurs within the PD (Figure 2A and B). The cluster includes the variants A314Y, S351Y, G354S, G356R and G375R. This PD cluster spans 20.3 Å, including residues from the transmembrane region S5, selectivity filter (SF), and the transmembrane region S6 that forms the inner vestibule. Inside the PD cluster, there is a sub-cluster in the SF between S351, G354, and G356 with distances of 5.85 Å between S351 and G354 and 3.66 Å between G354 and G356. The G354 and G356 residues are part of the well characterized ‘GYG’ signature sequence that confer the high K^+^ selectivity (31).

**Figure 2.**
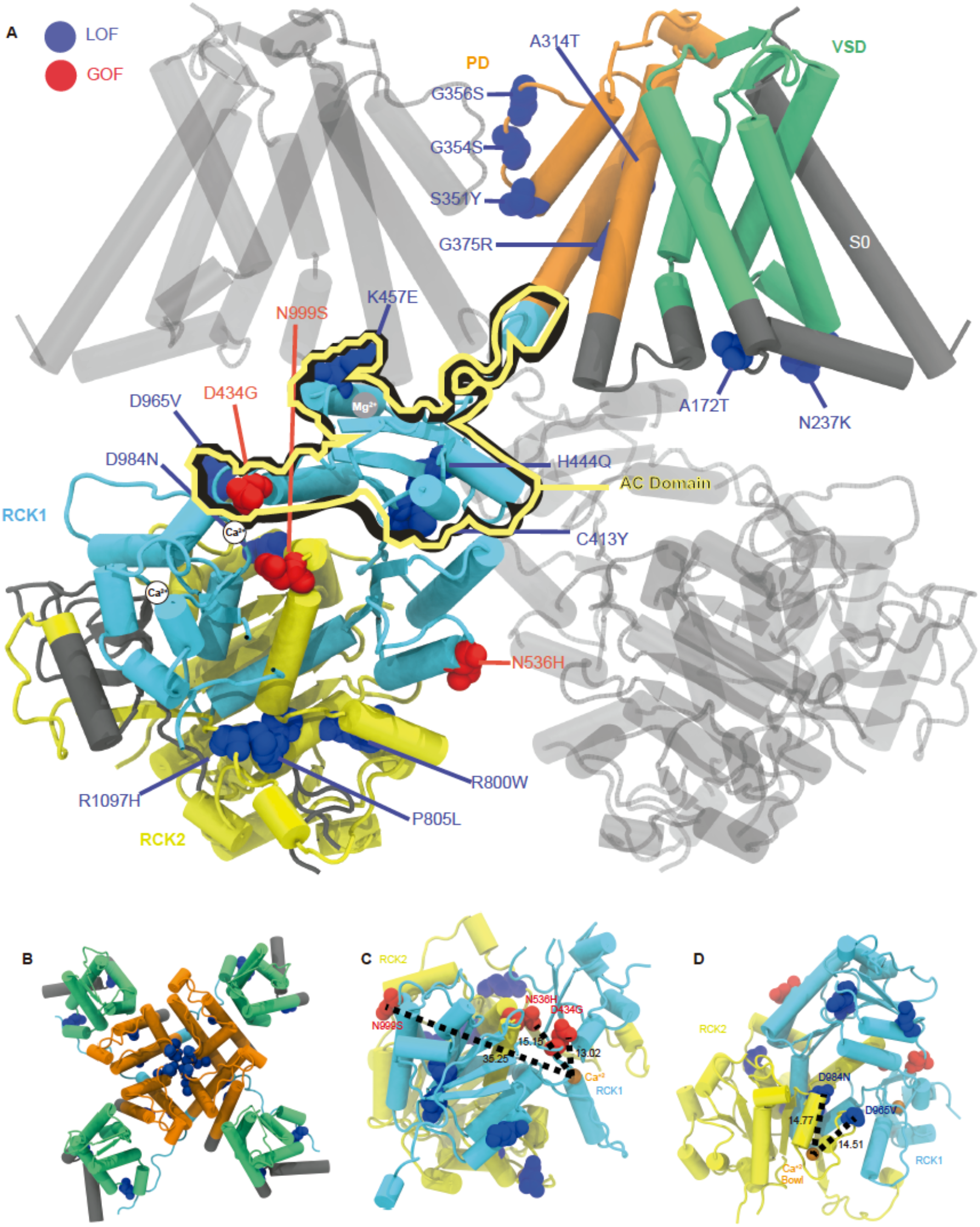
Clustering analysis of LOF and GOF HP variants. (A) Fifteen LOF (blue) and 3 GOF (red) variants mapped on the calcium bound BK channel. (B) Top view of the BK channel tetramer. (C) Magnification view of the gating ring showing the proximity of the GOF variants D434G and N536H to the RCK1 Ca^2+^ binding site. Dashed line measures distance in Å. (D) Magnification of the gating ring showing the proximity of the LOF variants D965V and D984N to the RCK2 Ca^2+^ bowl.

The second cluster of LOF variants occurs in the βA-αC (AC) region of the RCK1 domain (32) including the variants C413Y, H444Q, and K457E (Figure 2A and D). The variants R458ter and N449fs also fall in this subdomain but were not considered a part of the cluster as detailed above. The AC region cluster spans 22.6 Å, with variants C413Y and H444Q separated at 5.9 Å and H444Q and K457E separated by 14.7 Å. The relevance of the αβ helix of the AC region has been demonstrated in the past by mutagenesis, showing changes in the V_1/2_ when it is modified (32), similar to the phenotypes produced by LOF variants in this region (33). Interestingly, the well-characterized D434G GOF mutation could be considered part of this LOF cluster in terms of distance but not in terms of its GOF phenotype (27,32,34).

A pair of LOF variants, D965V and D984N, occur in a highly conserved subdomain of RCK2 containing a calcium binding site (Figure 2A) (16,35). D965V and D984N reside within 5 Å of the calcium bowl (3.4 Å and 4.5 Å, respectively), with a distance of 14.5 and 14.7 Å to the Ca^2+^ ion (Figure 2D). Another pair of LOF variants, R800W and P805L, also occur in RCK2. These residues are located 4.9 Å apart, at the end of S9 and in the S9-S10 linker (Figure 2A and D). While not considered a part of a cluster or pair, R1097H occurs close to the R800W/P805L pair, separated from the other two residues by 7.8 Å. This distance was not considered for the purpose of determining clustering here since the cutoff was 5 Å. However, values greater than 5 Å have been used in other clustering determination studies (36,37). Thus, the clustering of LOF HP variants showed a relationship between the structural location and the biophysical properties associated with the amino acid change.

Unlike LOF variants, the three GOF variants, D434G, N536H, and N999S, do not share an observable clustering pattern within 5 Å or localization within a single domain of the gating ring (Figure 2A). D434G and N536H are located in RCK1 (Figure 2A). The D434G residue is located in the AC region near the LOF cluster containing C413Y, H444Q, and K457E (38-41), but has the opposite electrophysiological phenotype producing a V_1/2_ shift to negative potentials (10,27,32,34). D434G and N536H are the closest variants to the Ca^2+^ binding site in the RCK1 with 13 and 15.1 Å, respectively (Figure 2C). Despite the proximity, D434G and N536H affect the gating properties in different ways, with D434G affecting the Ca^2+^ activation and N536H primarily affecting voltage activation (32,34,42). N999S, which is located in RCK2 35.2 Å from the Ca^2+^ site, also does not require Ca^2+^ activation to exert its effect over channel gating (41). N999S is located at the helix bend near the RCK1/RCK2 interface, 10 Å from D434G.

### Single Nucleotide Polymorphism (SNP) locations

While HP variants suggest deleterious effects on BK channel properties, SNP residues are a potential source of tolerated variability in the gating properties (8,25). The *KCNMA1* gene contains 537 non-synonymous SNPs (Supplemental Figure 2). Over half (298 SNPs) were located in resolved regions and could be mapped onto the BK channel structure. Of these, 142 SNPs were conservative residue changes based amino acid biochemical properties, and 156 were non-conservative. Figure 3 shows the 298 *KCNMA1* SNPs together with the GOF and LOF HP variants. Overall, there were many conservative SNPs at the N-terminus, associated with a high occurrence of synonymous variation (Supplemental Figure 2). Starting from the extracellular side, 52 SNPs occur in the unresolved N-terminus, and 33 SNPs are located in the glycine and serine repeats (Figure 3A, C). The intracellular loop between transmembrane segments S0 and S1 contains 38 SNPs, and 26 of these are non-conservative. The number of SNPs greatly decreases within the core alpha helices and transmembrane segments (Figure 3A and B). The transmembrane segments S3-S6 contain a total of 3 non-conservative SNPs, distributed between S3 and S5. In the gating ring, 39 SNPs occur after the end of RCK2 (Figure 3A and D), which includes 21 that are non-conservative. Finally, 154 SNPs are not shown due to their locations within unresolved structure regions. Thus, the BK channel N-terminus and gating ring are hot spots for SNP variability, in contrast to the VSD and PD.

**Figure 3.**
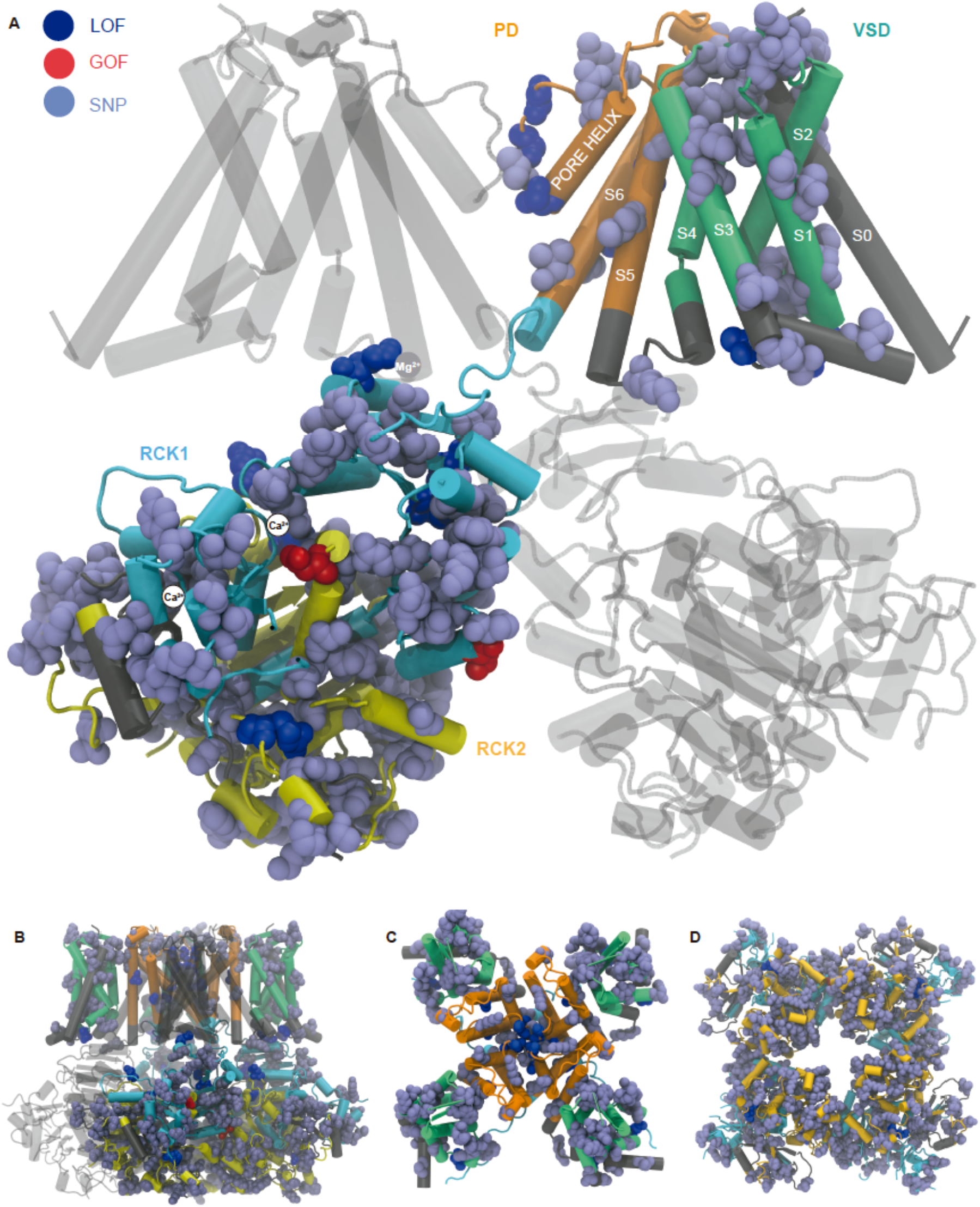
Distribution of non-synonymous SNPs by domain. (A) Localization of 60 SNPs (ice blue) alongside the GOF (red) and LOF (blue) HP variants in a two subunit-view configuration. (B) Side view tetramer representation. (C-D) Top and bottom (D) views. The Ca^2+^ binding sites at RCK1 and RCK2 (magenta circles) are located behind the area where SNPs R341, V370 and D371 (RCK1) and L447 and P451 (RCK2) are located.

Despite the presence of SNPs across the BK channel domains, several regions notably lack non-conservative SNPs. One is the pore (S5 to S6), a region critical for K^+^ permeation containing a cluster of LOF HP variants. Another region is the alpha beta helix that interfaces with the voltage-sensing domain in the AC region of RCK1, containing 3 LOF HP variants (Figure 3A). However, the opposite is observed in transmembrane segments S3 and S4, a region containing one non-conservative SNP but lacking HP variants (Figure 2A and 3A). Finally, the calcium bowl lacks SNPs, potentially due to the importance of this motif in the gating of the BK channel (43,44). Taken together, this analysis reveals domains with decreased non-conservative SNPs and HP variants. These domains are hypothesized to have lower tolerance to functional changes compared to domains with an increased number of SNPs, consistent with low SNP frequency in other proteins at modulatory sites (45).

### Sites of pharmacological modulation

BK channel gating properties are regulated by a variety of endogenous intracellular compounds or pharmacological agonists and antagonists. These modulators are relevant as potential therapeutics for *KCNMA1* channelopathy. To determine if HP variants were located near sites of modulation, binding sites and residues involved in modulation sites were localized with respect to HP variants within the BK channel structure. The inhibitors included in this analysis were charybdotoxin (ChTX) (46), iberiotoxin (IbTX) (47), loperamide (LOP) (48), paxilline (PAX) (49) and TEA (50,51). The agonists were BC5 (18), NS1619 (52), NS11201 (53) and Cym04 (52). The endogenous modulators were Heme (54), protons (H^+^) (55), lipids (56), carbon monoxide (CO) (57), kinases (58), and ethanol (EtOH) (59) (Figure 4A and B).

**Figure 4.**
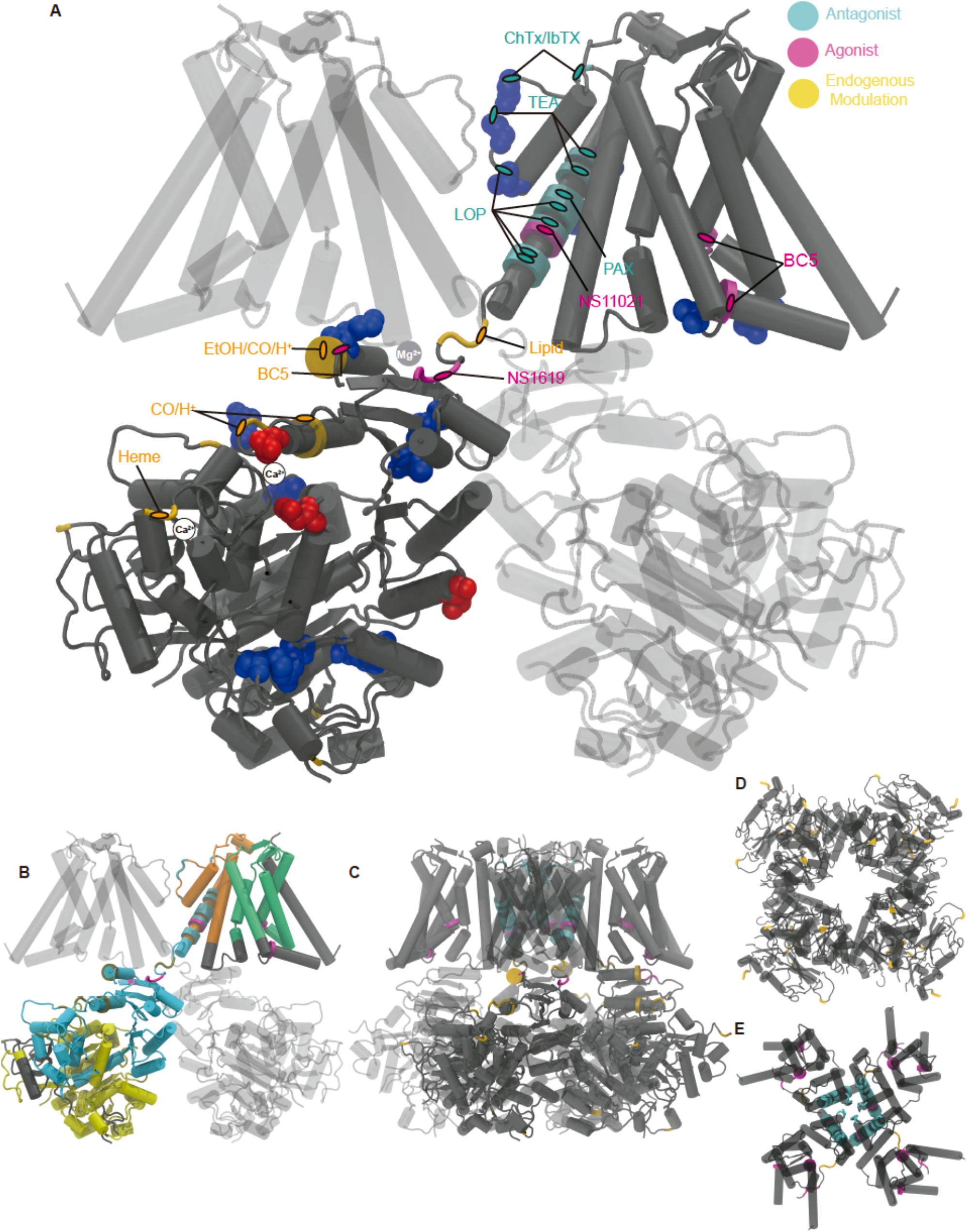
Sites of BK channel modulation and HP variants. (A) Charybdotoxin (ChTX), iberiotoxin (IbTX), loperamide (LOP), and paxilline (PAX) antagonist modulation sites are depicted in cyan. BC5, Cym04, NS1619, and NS11201 agonist modulation sites are shown in purple. Endogenous modulator sites (Heme, protons (H^+^), lipids, carbon monoxide (CO) and ethanol (EtOH)) are depicted in orange. 17 LOF (blue) and 3 GOF (red) HP variants are represented as globular residues. (B) Lateral two-subunit side view with domain representation: PD (orange), VSD (green), RCK1 (light blue) and RCK2 (yellow). (C-E) Four-fold symmetry representations from the side (C), bottom (D), and top (E).

The antagonists LOP, ChTX, IbTX and TEA (external binding site) cluster in the pore domain, acting as blockers in a range of 6.6 in the X axis and 8.3 Å in the Y axis. Interestingly, from the inner vestibule, the agonist NS11201 and antagonists LOP, TEA (internal binding site), and PAX cluster in S6 of the channel with 2.5 and 2.9 Å between residue binding sites (Figure 4A and E). NS11201 shares part of the modulation area with LOP, despite its opposite agonist effect over the BK channel activation. This S6 modulation cluster overlaps with the LOF HP variants S351Y, G354S, and G356S, and G375R, a mutation producing small LOF effects on conductance but large GOF effects on gating (60). The ChTX and IbTX binding sites are located in the extracellular loop that connects the pore and S6, composed by site 1 in the selectivity filter and other residues in the turret that stabilize block. No HP variants are currently found in this specific area (Figure 4E).

In the case of the endogenous intracellular modulators, residues involved in EtOH (activator), CO (activator), lipid (activator), and H^+^ (activator) modulation cluster at the start of RCK1, at distances of 4.2 and 9.5 Å respectively. Overlapping residues are found between CO, H^+^, EtOH, and the LOF variant K457E (Figure 4A and D). Additionally, the sites overlap at 3.8 Å of distance with the BC5 agonist at the start of RCK1.

Marking sites of kinase modulation, 30 residues were detected as phospho-sites by mass spectrometry from rat brain tissue (58). Only 4 of these residues were located within resolved areas of the structure: S582 (occurring at the end of S7 in RCK1), S850 (between S9 and S10 in RCK2), T1065 (end of RCK2 after S10), and S1123 (after RCK2). No clustering was observed between any of these 4 phospho-sites and HP variants. However, due to the low number of phospho-sites within resolved regions, no conclusions are currently suggested. Generally, the analysis of modulator and binding sites showed a low number of proximal or overlapping HP mutations. One exception was the pore domain, where the LOF variants and several pharmacological inhibitors have effect over BK channel activity in the same direction.

### *KCNMA1* HP variants producing putative truncations

Four nonsense HP variants are predicted to produce truncated BK channel proteins: R458Ter, R640Ter, R830Ter, and R880Ter (61) (ClinVar and ALM unpublished data). Assuming proper folding of the channel with the deleted residues, these putative truncations were schematized to determine the relationships between structural boundaries and BK channel terminations (Figure 5A-D). The representations were then compared with two experimentally truncated channels: Slo1C and Slo1ΔRCK2. Slo1C terminates in the linker between S6 and RCK1 at position 342 and is missing the whole gating ring. Slo1C is characterized as a LOF mutation by electrophysiology, affecting the allosteric interactions in the BK channel gating mechanism (17). Slo1ΔRCK2 terminates after RCK1 at position 632. It has not been studied functionally, but clearly delineates the RCK domain boundaries in the gating ring domain.

**Figure 5.**
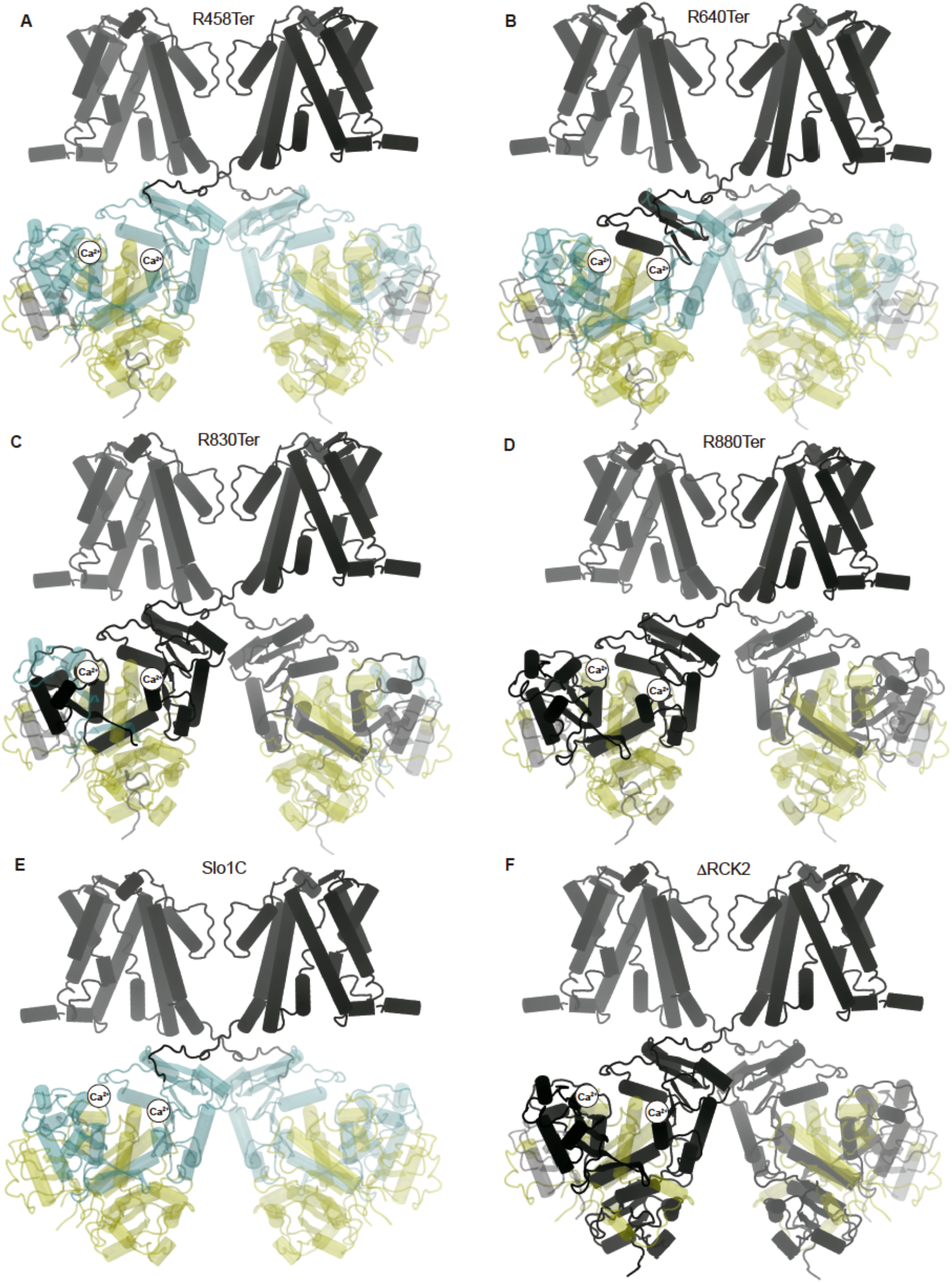
HP patient variants creating premature stop codons. (A) R458Ter translation terminates at the end of the AC domain. (B) R640Ter terminates in RCK1 before the alternative splice site 1. (C) The R830Ter and (D) R880Ter terminations occur before and after S9, respectively, prior to the calcium bowl located in the RCK2. (E) Slo1C truncated BK channel. Slo1C also includes a Kv1.4 tail of either 74 or 11 amino acid residues (17), which is not shown in the structural representation. (D) SloΔRCK2 is an in silico truncated BK channel for reference, showing the boundaries of RCK1 and RCK2. Black and transparent blue/yellow denote the portions of channel before and after the putative truncations, respectively.

The R458Ter variant has a LOF phenotype with the putative truncation occurring in the AC region of RCK1 (Figure 5A) (62). The R640Ter variant creates the putative truncation before the alternative splice site 1 in RCK1 (Figure 5B) (61). The R830Ter variant creates a putative termination before S9 at the start of RCK2 (Figure 5C). All three are classified as pathogenic by ClinVar. Variant R880Ter is located within RCK2 with the truncation occurring after S9 and before the calcium bowl (Figure 5D) and is classified as VUS. Only R458Ter has direct electrophysiological evidence for a LOF classification. The three other truncations (R640Ter, R830Ter, and R880Ter) are functionally unstudied at present.

Deletions within the BK channel gating ring could create a LOF BK channel due to loss of the calcium binding sites (63). The Slo1C truncation occurs at the end of the tetramerization domain after S6 and before RCK1 at position 342 (Figure 5E). Slo1C and R458Ter have a similar termination positioning and result in comparable electrophysiological properties (8,17). However, patients are heterozygous for these variants, and these truncations have the potential to produce more complex BK channel alterations. Truncations could also theoretically produce a GOF effect, if some structural or allosteric restrictions from the gating ring are removed. In the case of R830Ter and R880Ter, which along with SloΔRCK2 have not been characterized electrophysiologically, one Ca^2+^ site is deleted. However, if functional BK channels are formed from these variants, it is more likely that the final effect of a putative truncation at these positions would result in a LOF in trafficking or gating properties (17).

### Assessment of *KCNMA1* HP variants by pathogenicity algorithms

Variant pathogenicity is commonly initially evaluated by tools available in ClinVar, Ensembl, Varsome, and others (19). The most widely used algorithms are Swift (64), Polyphen (65), CADD/PHRED (66), MetalLR (67), M-CAP (68), Mutpred2 (69) and REVEL (70). These genetic algorithms are divided into independent or ensemble predictors (19). The independent predictors (SIFT, Polyphen, and Mutpred2) analyze the tolerance of the residue to change by statistical or evolutionary sequence comparison methods (64,65,69). The ensemble predictors (CADD/PHRED, MetaLR, M-CAP, and REVEL) use combination of independent predictors through machine learning methods (67,68,70,71). While these genetic pathogenicity algorithms appear to perform well for missense variants of less complex structural proteins, these tools may not be adequate to assess mutations in genes encoding ion channels (21). For example, Polyphen and Mutpred incorporate protein structural information by amino acid residue accessibility or effects over protein structure, respectively. Each uses trained datasets that do not necessarily represent ion channel structural domains such as the voltage sensor, inactivation motifs, ion binding sites, and allosteric pathways. Neither incorporate structures from specific conformational states, such as open versus closed channels. Therefore, existing individual and ensemble tools may be inadequate to assess pathogenicity associated with some genes (19,21).

To address this, we evaluated the performance of SIFT, Polyphen, CADD/PHRED, MetaLR, M-CAP, Mutpred and REVEL with nonsynonymous *KCNMA1* variants (Figure 6). Frameshift, deletion, or alternative exon variants were not included. Four parameters were assessed-specificity, sensitivity, accuracy, and precision (see Methods) using 16 pathogenic variants confirmed by BK channel activity and 4 benign variants with no effect on BK channel activity. Of these algorithms, CADD/PHRED, MetaLR, Mutpred and REVEL showed the best predictive capabilities in each parameter (Figure 6A), with average performance metrics for the four parameters of 0.78, 0.80, 0.87, and 0.88 respectively (Figure 6B). Polyphen also showed better performance than CADD/PHRED and MetaLR (0.81). However, because it is incorporated into the ensemble algorithm REVEL, Polyphen was left out of this analysis. M-CAP was also not used for further analysis because one of the individual score components (specificity) was lower than the other algorithms.

**Figure 6.**
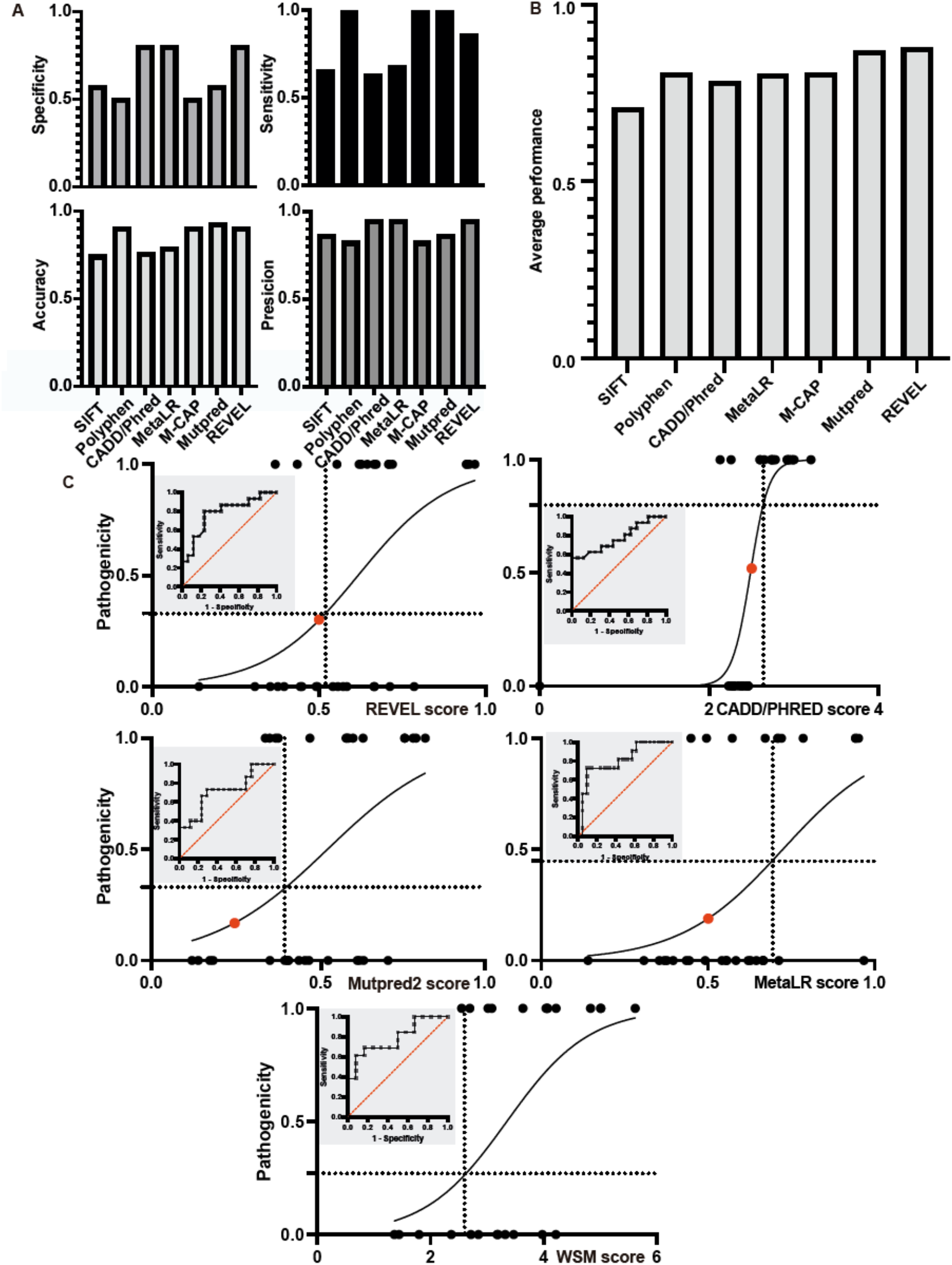
Evaluation of individual and score-ensemble algorithms for *KCNMA1* pathogenic variants. (A) Accuracy, precision, specificity, and sensitivity for *KCNMA1* pathogenic and benign HP variants (left) and summary score (right). (B) Pathogenicity score plots with logistic regression curves generated from the training data set for the four algorithms showing the best average performance. Dotted lines denote the experimentally-derived cutoff values from the training data set (CADD/Phred, 2.7; MetaLR, 0.71; Mutpred, 0.43; REVEL, 0.51; and WSM, 2.6), corresponding to true positives (65-90%) and false positives (25-45%). The red dots indicate the standard cutoffs from ClinVar (CADD/Phred, 2.4; MetaLR, 0.5; Mutpred, 0.25; REVEL, 0.5). Insets: Receiver operating characteristic (ROC) curve showing true positives (pathogenic variants detected as such; sensitivity) against the false positives (non-pathogenic variants detected as such; 1-specificity). The linear regression for a perfect correlation is in red.

After determining the performance with each algorithm, logistic regression was used to estimate the probability of a pathogenic (1) versus non-pathogenic (0) categorization for the HP variants confirmed by BK channel recordings in the training data set (Figure 6C). Receiver operating characteristic (ROC) curves were constructed, which assess the diagnostic ability of the binary pathogenicity designations by plotting the true positives (pathogenic variants detected as such) against the false positives (non-pathogenic variants detected as such, Figure 6C insets). This probability assessing whether a randomly chosen pathogenic variant will be ranked higher than a randomly chosen non-pathogenic variant was used to determine the best cutoffs for each algorithm (see Methods). Surprisingly, the analysis revealed that using general ClinVar cutoffs was associated with a high false positive rate (Supplemental Table 1B and Figure 6C). The result demonstrates a low discrimination power of the common genetic algorithms between pathogenic and non-pathogenic variants when used with general cutoffs on the known *KCNMA1* dataset.

### Development of a structurally-integrated algorithm for *KCNMA1*

To assess the performance of the existing algorithms using structural information from this study, we developed a building block strategy that combines the pathogenicity algorithms and structural parameters. The first step consists of a weighted summation module (WSM) that incorporates the sum of the best-performing individual algorithms REVEL (R), Mutpred (MP), MetaLR (MLR), and CADD/PHRED (CP). The WSM value was then adjusted by the capability of each algorithm to detect true positive and negative variants. Each individual score was multiplied by the fraction of true positives (pathogenic variants) and divided by the fraction of false positive (false pathogenic variants) that was able to detect and incorporated in the model according to equation 5 (Supplemental Table 1):

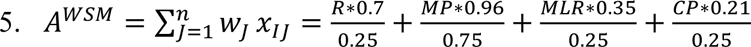

w: weighted factor and x: each individual algorithm (R, MP, MLR, and CP). From the ROC curve, the cutoff associated with a higher true positive detection, as well as a decrease in false positives, was obtained with the WSM (Figure 6C). For example, at a cutoff of 2.6, the performance of WSM is better than REVEL in the true positive rate determination (88 versus 85%).

The second step in the KMS generates a structural component (SC) that extends beyond REVEL and Mutpred, two algorithms which incorporate residue conservation, properties, and secondary structure. The SC includes the number of interactions (NI), number of residues (NR), and associated energy of interaction (AE) averaged from BK channel structures in the absence and presence of Ca^2+^ (PDB:6V3G and 6V38) (16,22), making the general assumptions that functionally important residues would: 1) be located within 5 Å of other residues, 2) interact with other residues via hydrogen bonding, π-π stacking, π cation, salt bridge, Van Der Waals or disulphide bonds, 3) have high energetics of interaction. The first SC parameter is the average number of residues (NR) at a 5 Å distance from the residue of interest that represents a cluster and possible interaction. The second parameter was the average number of interactions (NI) that represent structural stability, and the third parameter was the average energetic value of those interactions (AE). These parameters were combined in the structural component (equation 6; Supplemental Table 2):

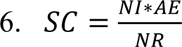

For the residues not resolved in the structure or with NI values of 0, we used minimal NR and/or NI values of 1 and the average energy of a hydrogen bond interaction of 18 KJ/mol as AE for the ND and 1 for NI values equal 0 (Supplemental Table 2). These minimal values were used to facilitate evaluation of the complete set of *KCNMA1* HP variants, avoiding indeterminates created by a zero term.

The KMS formula is:

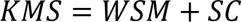

The cutoff for significant KMS scores was generated on a training data set of pathogenic and non-pathogenic variants (Figure 7A). The cutoffs calculated for WSM (2.6) and KMS (2.5) showed a similar true positive detection probability (89%) with a reduced false positive rate of 75% compared to the WSM alone 50%, compared from the ROC curves at 0.4 of 1-specificity for both scores. Incorporation of the structural parameters in the KMS algorithm also results in the highest area under the ROC curve (AUC) among the different algorithms (0.85), compared to both WSM and REVEL (0.78 for each), suggesting a better cumulative performance compared to the available ClinVar pathogenicity algorithms.

**Figure 7.**
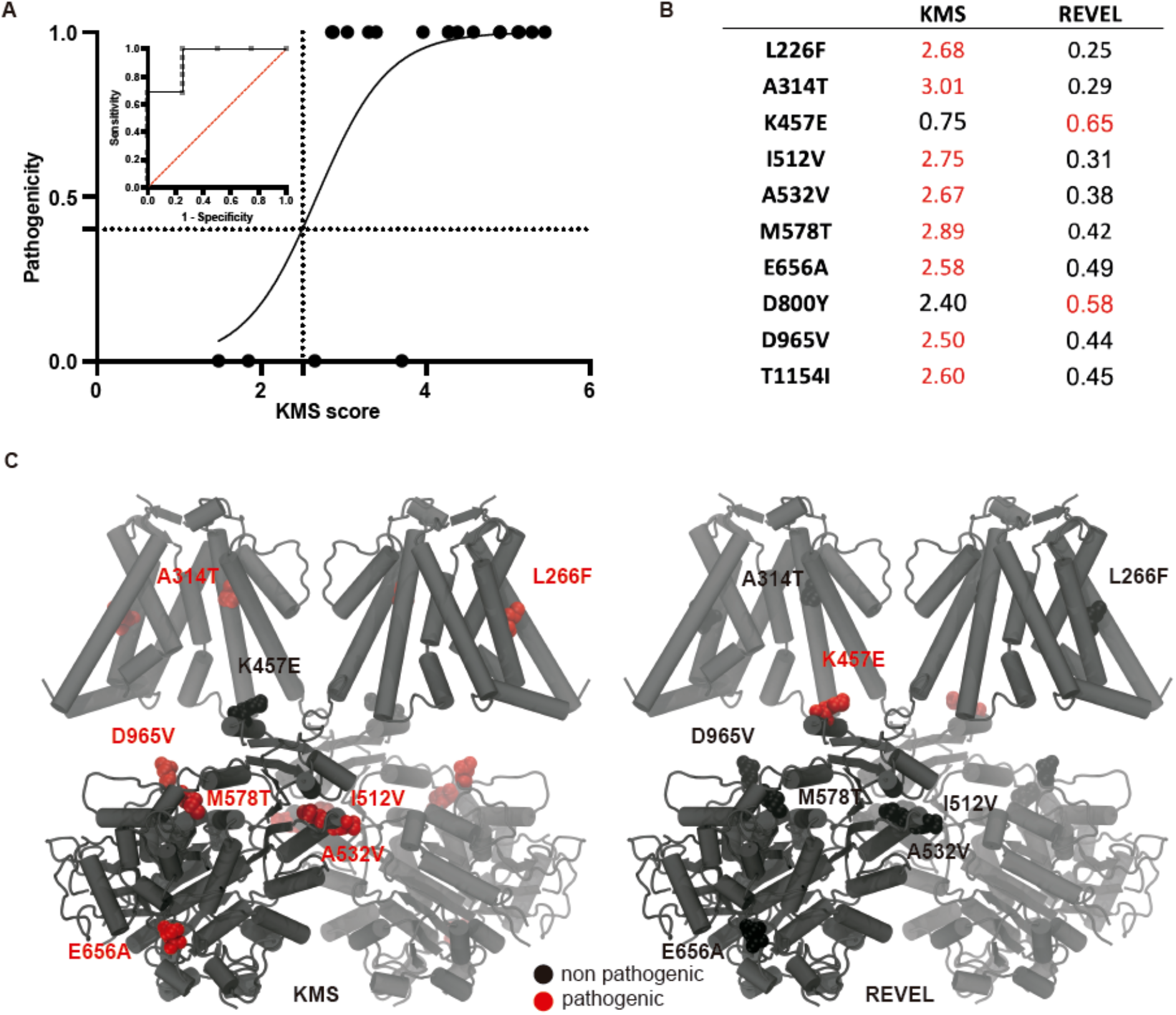
*KCNMA1* Meta Score (KMS). (A) Pathogenicity score plot with logistic regression curve generated from the training data set. Dotted lines denote the cutoff of 2.5, corresponding to a true positive rate of 88% (sensitivity) and 45% false positive rate (1-specificity). Inset: Receiver operating characteristic (ROC) curve with linear regression for perfect correlation (red). (B) KMS and REVEL algorithm scores for VUS residues with conflicting predictions; scores above the cutoffs are red (assessed as likely pathogenic) and below the cutoffs are black (assessed likely non-pathogenic). (C) Ca^2+^-bound human BK channel cryo-EM. D800Y and T1154I are not depicted because they are located in unresolved structural regions.

### Electrophysiological evaluation of HP variants delineated by KMS score

To evaluate the KMS algorithm on VUS *KCNMA1* HP variants, we took two approaches: 1) comparing KMS scores to REVEL, a widely used algorithm showing the best overall individual performance but lacking BK channel-specific substantiation, and 2) validating a subset of VUS residues that showed conflicting predictions with respect to BK channel activity using electrophysiology. KMS showed a 12% difference from REVEL (Figure 7, Supplemental Table 1B), suggesting that using a trained ensemble algorithm results in a small performance improvement associated with identifying patient mutations that change homomeric BK channel activity. Ten variants were identified with differing KMS and REVEL outcomes (Figure 7B). Eight VUS variants were assessed as potentially pathogenic by KMS but not by REVEL. These 8 VUS had KMS scores above than the KMS cutoff, and REVEL scores that were below than the REVEL cutoff (KMS^+^; REVEL^-^): L226F, A314T, I512V, A532V, M578T, E656A, D965V, and T1154I. Two VUS were assessed as likely pathogenic by REVEL but potentially non-pathogenic with KMS (KMS^-^; REVEL^+^; Figure 7B-C): K457E and D800Y. Four of these mutations were tested functionally in this study using electrophysiology (M578T, E656A, D800Y, and D965V). These four variants were evaluated alongside two additional novel variants (G20D and R1097H) and eight already classified HP variants that were previously studied under different conditions (1,8). Collectively, Figure 8 reports the functional effects on BK current properties for a total of 13 HP variants under identical conditions (10 μM Ca^2+^). Evidence for pathogenicity at the BK channel level was considered based on a change in the voltage-dependence of BK current activation (V_1/2_), activation kinetics (τ_act_), or deactivation kinetics (τ_deact_; Figure 8) compared to wildtype (WT).

**Figure 8.**
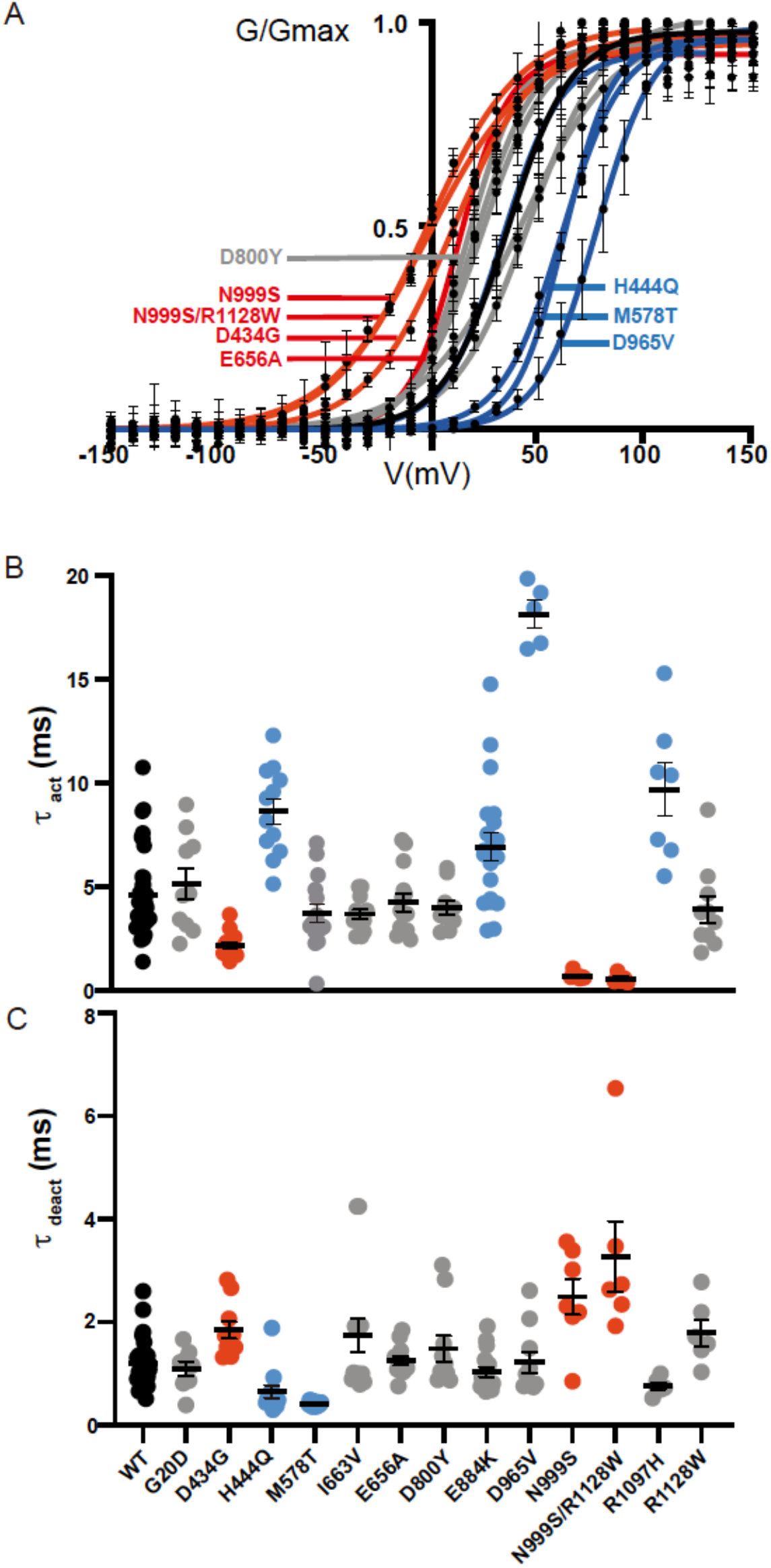
Electrophysiological characterization of *KCNMA1* HP variants. (A) Conductance versus voltage relationship curves of the WT and 13 patient variants in the human BK channel. Inside-out patch-clamp recordings were conducted in a physiological K^+^ gradient and 10 μM intracellular Ca^2+^ conditions. Data average ± SEM are plotted with Boltzmann fits. V_1/2_ values (mV): WT (black): 29 ± 3 (N=27). Variants with hyperpolarizing shifts in the V_1/2_ (red): D434G 7 ± 4 (N=12, P=0.0001), E656A 12 ± 5 (N=13, P=0.017), N999S -15 ± 3 (N=14, P<0.0001), and N999S/R1128W -8 ± 7 (N=7, P<0.0001). Variants with depolarizing shifts in the V_1/2_: H444Q 52 ± 5 (N=13, P=0.0006), M578T 65 ± 5 (N=19, P<0.0001), and D965V 69 ± 6 (N=10, P<0.0001). VUS/no effect (gray): G20D 20 ± 6 (N=9, P=0.76), I663V 15 ± 6 (N=11, P=0.13), D800Y 21 ± 6 (N=10, P=0.81), E884K 42 ± 5 (N=20, P=0.1007), R1097H 21 ± 5 (N=14, P=0.69), and R1128W 51 ± 8 (N=5, P=0.068). (B) Activation kinetics at +30mV, a voltage relevant during neuronal activity. WT 4.48 ± 0.03. Variants with activation kinetics slower than WT: H444Q 8.52 ± 0.62 (P<0.0001), E884K 6.80 ± 0.67 (P=0.0003), D965V 18.01 ± 0.66 (P<0.0001) and R1097H 9.56 ± 1.29 (P<0.0001). Variants with activation kinetics faster than WT: D434G 2.07 ± 0.12 (P=0.0003), N999S 0.60 ± 0.06 (P<0.0001) and N999S/R1128W 0.48 ± 0.099 (P=0.0003). Variants with no effect on activation kinetics: G20D 5.03 ± 0.73 (P=0.99), M578T 3.60 ± 0.45 (P=0.81), I663V 3.57 ± 0.22 (P=0.83), E656A 4.15 ± 0.43 (P=0.99), D800Y 3.88 ± 0.33 (P=0.99) and R1128W 3.80 ± 0.64 (P=0.98) (C) Deactivation kinetics at -70mV, approximating an average neuronal resting membrane potential. WT 1.20 ± 0.23. Variants with deactivation kinetics slower than WT: H444Q 0.65 ± 0.17 (P=0.0013) and M578T 0.42 ± 0.21 (P=0.0044). Variants with deactivation kinetics faster than WT: D434G 1.85 ± 0.23 (P=0.044), N999S 2.49 ± 0.27 (P=0.0001) and N999S/R1128W 3.27 ± 0.28 (P<0.0001). Variants with no effect on deactivation: G20D 1.1 ± 0.26 (P=0.99), I663V 1.74 ± 0.25 (P=0.17), E656A 1.25 ± 0.23 (P=0.99), D800Y 1.48 ± 0.24 (P=0.95), E884K 1.03 ± 0.16 (P=0.83), D965V 1.21 ± 0.19 (P=0.99), R1097H 0.78 ± 0.30 (P=0.80) and R1128W 1.89 ± 0.31 (P=0.41).For B-C, individual data points ± SEM are plotted. P values correspond to Bonferroni post-hoc test from two-way ANOVAs.

The mutations D965V and M578T both showed a strong LOF phenotype for BK currents (Figure 8A-C). This LOF effect was large when compared to other LOF variants reported in the literature, such as H444Q (11,38). Neither D965V nor M578T variants are located within a cluster in the structure; however, these residues were chosen for functional analysis because both are near, or part of, the calcium activation structures. M578T (M513 in the cryo-EM structure) is part of the Ca^2+^ binding site in RCK1 (formed by D429and D436; Figure 7C). M578 was previously mutated to isoleucine in another study, showing a disruption in the Ca^2+^ activation and a LOF phenotype (43,72). M578T produces a similar LOF phenotype as M578I with a shift of +34 mV (Figure 8A). The D965 residue is near to the calcium bowl in the RCK2 (DDDDDP<u>D</u>) (73). Previously, the D965A mutation showed no difference from WT between 0 and 100 μM Ca^2+^ (43,72). However, in this study the HP variant D965V showed a LOF phenotype with a + 40 mV shift in the V_1/2_ (Figure 8A). Under these conditions both variants alter BK channel properties; however, D965V produces a stronger LOF phenotype than M578T.

The E656A variant was also assessed as potentially pathogenic by KMS but not by REVEL (Figure 7B-C). The residue E656A is in the region between S8 and S9, pointed to the gating ring (Figure 7C). E656A was previously tested, showing no differences compared to WT BK currents in the absence or presence of the accessory subunit β4 (74). BK current recordings in this study showed a -17 mV hyperpolarizing shift in the V_1/2_ (Figure 8A) but no concomitant change in kinetics (Figure 8B-C). Under these testing conditions the change in BK currents was consistent with the KMS assessment, but whether the variant could be classified accurately as GOF requires additional investigation to resolve the inconsistent functional data.

The D800Y variant, assessed as likely pathogenic by REVEL but not by KMS (Figure 7B), did not show any difference in BK current properties compared to WT (Figure 8). This finding is also in accordance with KMS over REVEL. However, D800Y is one of the HP variants that cannot be localized in the structure because it lies within an unresolved region. In this situation, KMS will perform at the WSM detection limit using an arbitrary value for the structural component and does not indicate predictive capability from structural integration in the KMS. However, as with all putative benign residues, the effects on expression, protein-protein interactions, and other conditions, remain to be tested.

One other variant (K457E) was also assessed as potentially non-pathogenic by KMS, but likely pathogenic by REVEL. K457 is in the αB helix of RCK1 (Figure 2A) and interacts with Phosphatidylinositol 4,5-bisphosphate (PIP2) to promote BK channel activation (75, 95). Although K457E was not tested by electrophysiology in this study, it was tested in a previous experiment. Neutralization of the K457 positive charge (K392N) decreased PIP2 sensitivity and shifted the V_1/2_ to more depolarized potentials (75). K457 is also implicated in interactions with β subunits, and K457E in tandem with an additional mutation R393D decreased activation of the α-subunit and the β subunit-mediated increase in Ca^2+^ sensitivity (76,77). These combined results suggest that K457E is likely a LOF variant (78), and is potentially incorrectly assessed by KMS at the BK channel level.

Five additional HP variants L226F, A314T, I512V, A532V, and T1154I were identified as potentially pathogenic by KMS score, but not REVEL (Figure 7B). The effects of these mutations on BK currents were not tested in this study. Two additional novel VUS that did not have conflicting KMS and REVEL pathogenicity assessment were also studied by electrophysiology: G20D and R1097H. G20D, located in the N-terminus but not resolved in the cryo-EM, was assessed as likely non-pathogenic with KMS. Consistent with this, BK current recordings showed no difference compared to WT (Figure 8A-C). R1097H, located in RCK2 near the R800W/P805L LOF pair, was assessed as potentially pathogenic by KMS. In these conditions, BK current activation was 2.1 times slower compared to WT; however, the V_1/2_ and deactivation were not different (Figure 8A-C). R1097H showed greater LOF changes at lower Ca^2+^ (1 μM), with a +44 mV in the V_1/2_ (WT n=16 and R1097H, n=8, p<0.0001), 7.4 times slower activation (at +160mV; p<0.0001), but no change in deactivation. The remaining variants studied by electrophysiology in Figure 8 were a subset of variants used in the training data set.

## Discussion

In this work, we analyzed the subset of nonsynonymous *KCNMA1* patient-associated variants found in constitutive exons with respect to their locations and pathogenicity. Fifty-three residues and 4 putative truncations were resolved within the open and closed BK channel cryo-EM structures (16,79,80) and mapped with respect to channel domains, functional effects (GOF versus LOF), and overlap with sites of endogenous and pharmacological BK channel modulation and natural genetic variation (SNPs). The combined picture that emerges from the currently reported set of HP variants suggests that disease association identifies new residues within the VSD, pore, and gating ring that modulate BK channel activation. Surprisingly, no variants occur directly in the gating charges, Ca^2+^ binding sites, or other well-studied allosteric sites. Few HP variants correspond to residues identified as important in prior gating studies, suggesting their disease association has the potential to inform new residues controlling critical gating parameters. This observation was most notable among the three GOF HP variants producing the strongest disease correlation (8). The strong association with GEPD (generalized epilepsy and paroxysmal dyskinesia) predicted mechanistically similar channel-level alterations between D434G, N536H, and N999S. However, no clustering or structural similarities between these variants was found. How these variants produce similar GOF BK channel phenotypes, despite their disparate locations, differing Ca^2+^-dependent (D434G; 32) and independent (N536H and N999S; 41, 42) mechanisms, and in the heterozygous configuration, remains to be determined. Other variants with more limited GOF gating effects (G375R and E656A) do not share the same patient symptomology (60,74)..

In contrast to the rarity of GOF variants and lack of structural themes at the channel level, most LOF variants (80%) localize within 5 to 22 Angstroms of each other (Figure 2). Predictably, most are found within the pore region, which is essential for K^+^ selectivity and permeation and largely devoid of non-conservative SNPs. The variants mapping to pore residues overlap with a major region of pharmacological inhibition in the BK channel. Other LOF variants cluster within the AC domain of RCK1, a second domain emerging as an identifiable cluster of functionally consequential residues. This region also lacks non-conservative SNPs. Interestingly, modulatory sites (H^+^, CO and EtOH) in the AC region or RCK1 are close to pathogenic HP variants. However, the effects of the HP variants over these modulations, or vice versa, have not yet been evaluated. Interestingly, individuals harboring D434G variants have been reported to experience alcohol-induced PNKD (10). Thus, localizing variants with common mechanistic channel properties (GOF versus LOF) with respect to BK channel domains, sites of regulation, or distinguishing symptoms in the individuals harboring these variants provides a new basis to analyze genotype-phenotype correlations.

Pathogenicity was assessed using multiple established algorithms, incorporating new cutoffs derived from functional studies within a training dataset, and integrating these analyses with structure-based parameters into an ensemble KMS algorithm to prioritize several VUS for functional studies. Despite the low number of HP variants affecting BK channel gating that have been functionally confirmed by electrophysiology in the literature, the ensemble KMS algorithm showed some initial potential augment the standard genetic-based algorithms beyond their individual performances. Incorporation of structural features (residue interactions and state-dependence) and establishing cutoffs from a training dataset of confirmed BK channel-level pathogenic HP variants using KMS resulted in a re-consideration of three REVEL-predicted non-pathogenic VUS residues as potentially pathogenic (M578T, E656A, and D965V), and one REVEL-predicted pathogenic residue (D800Y) as potentially non-pathogenic. KMS performed well with residues that are close to regulatory sites such as the calcium bowl (M578T and D965V), which was confirmed by electrophysiological data. KMS also delineated HP variants in regions not resolved by the cryo-EM structure (G20D and D800Y). However, since this observation lacks structural information, the WSM score also assessed such residues as correctly according to the BK current recordings. On the other hand, this algorithm failed to accurately categorize K457E, a residue of known function in a structurally-resolved region, identifying that KMS predictive capabilities will need to be increased as more disease-delineated variants are functionally tested beyond the 16 used for training this dataset, more conditions are tested, new structural and allosteric components are available, and novel functional modules capable of assessing heteromeric channels and expression are developed. Moreover, the biophysical characterization in this study only tests a single Ca^2+^ condition, providing initial evidence for changes in the voltage-dependence of activation or kinetics but not the mechanisms behind them. The modularity of KMS was designed to evolve with improved genetic analysis and structural resolution, which only involves replacing terms in the equation with new modules as the component algorithms are improved.

Pathogenicity analysis in other ion channels has required similar approaches (21,82,83), but despite the evolution, there are global and gene-specific limitations. Those limitations for KMS in its current form are based on the structural components and how they are incorporated. The assumption of minimal values in the number of interactions (NI), number of residues (NR), and average energy (AE) is only true if the variant residues are located within a structured region of the channel that is not currently resolved in the cryo-EM structures. NI and NR may also underemphasize functionally important pair interactions. This could be partially solved with a term that accounts for the folding impact of a given mutation, helping to discard neighbor interactions. Similarly, AE may underemphasize important allosteric pathways and sub-states, since it only reflects the Ca^2+^-dependent open and closed states. Additionally, residues within other lipid, ion, and protein binding sites are difficult to incorporate into the equation in its present form without affecting the data dichotomization and cutoff determination. This affects variants N237K (part of the Mg^2+^ binding site) (84,85) and K457E (PIP2 binding site). In addition, the limited KMS training data set composed of 16 pathogenic and 4 non-pathogenic residues, could underestimate the performance of some scores, penalizing them in the WSM and affecting the logistic regression dichotomization. Other limitations relevant to BK channel pathogenicity prediction are not restricted to KMS (ie they are also present in CADD/PHRED, MetaLR, Mutpred and REVEL), such as the inability to predict transcript abundance or channel expression defects. However, KMS could potentially incorporate an expression-based module in future iterations, derived from in silico sequence prediction or from direct functional expression studies.

Another consideration is the relevant composition of BK channels in vivo, which KMS and other pathogenicity algorithms cannot currently assess. Most *KCNMA1* HP variants are heterozygous alleles (97%). This scenario could create heterotetramerization between WT and mutant subunits, potentially decreasing or altering the pathogenic phenotype within the cellular context, depending on the biophysical mechanism and its degree of dominance. This was observed for BK^R207Q^ GOF channels co-expressed with BK^WT^ (86) as well as in the A172T, A314T, G375R, N536H and N999S (1,39,42,60). Interestingly, homotetrameric G375R was reported to decrease single channel conductance by 40%, consistent with the decreased currents observed at its initial report (38,60). However, in a more detailed analysis, both homotetramers and G375R co-expressed with WT subunits showed a net GOF effect via a massive hyperpolarizing gating shift (60). At present, the stoichiometry, properties, and expression of heterotetrameric channels cannot be predicted and must be addressed through functional testing.

Lastly, KMS and other algorithms are binary. The score magnitude is not associated with the degree of pathogenicity and will not predict results in the context of heterozygous alleles in vivo. Most ion channelopathies face similar challenges as those encountered in *KCNMA1* channelopathy (87). Several studies affirm that these aforementioned tools fail to accurately assess pathogenicity for certain classes of proteins, including some ion channels (88-92). The detection pitfalls mainly result from the use of a specific threshold generated from true positive (sensitivity) versus false positive (1-specificity) receiver operating characteristic (ROC) curves, using non-optimal datasets to generate binary classifications of pathogenic (1) and non-pathogenic (0) (19,88). Individual algorithms that use genetic or amino acid sequence datasets miss unique motifs in ion channels like regulatory elements, ‘non-shaker’ voltage sensing domains and other gating elements (30), regulatory binding sites (93), and residues relevant for the interface interactions in multimeric complexes (94). This is most relevant for ion channels defined by a single family member or those with low homology regions across genomes. Meanwhile the ensemble algorithms do not include a threshold correction, leading to the loss of control over Type I error (rejection of null when true) and the wrong identification of non-pathogenic variants as pathogenic (Type II error, failure to reject null when false). Together these errors contribute to identifying pathogenic variants as non-pathogenic (19). More recently, the sodium channel channelopathies (as with other genetic diseases) have addressed these issues recalculating the pathogenicity threshold of different scores using data from *SCN*1A, 2A and 8A-specific patients, improving the disease driven variants (21). Thus, the development of new algorithms, or improvement of standard ones, has the potential to push a better predictor performance toward faster genotype-phenotype categorizations. Ideally, computational pathogenicity assessment would incorporate multi-modal genetic, structural, molecular dynamic, channel function, and neuronal evidence into an ensemble predictor tool. This type of higher-order analysis is increasingly being developed in a gene- and disease-specific context (21,88-92).

## Acknowledgements

We gratefully acknowledge patients, families, and physicians for their participation in the KCNMA1 Channelopathy International Advocacy Foundation (KCIAF) registry questionnaire or providing direct submission information, and Alyssa Mendel and the Coordination of Rare Diseases at Sanford (CoRDS) program for providing patient registry data collation and access.

## Author contribution

HJM: research conception and data collection, statistical analysis design and execution, and manuscript writing. KT: research conception, data collection, and manuscript writing. ALM: research conception and data analysis, statistical analysis design, and manuscript writing.

