## Supplementary figures and images for "Structural mapping of patient-associated *KCNMA1* gene variants"

### Supplemental Figure 1

#### Supplemental Figure 1

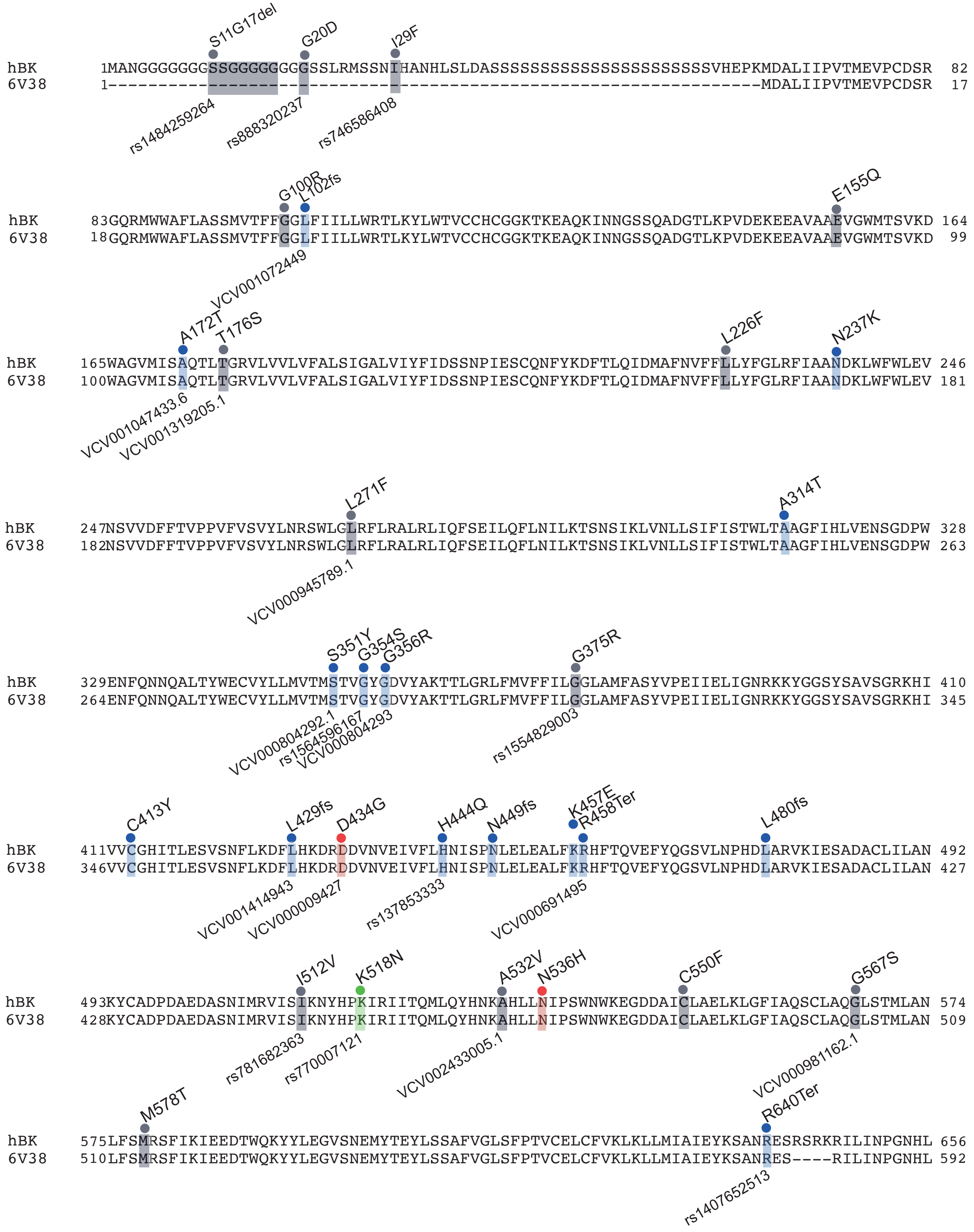

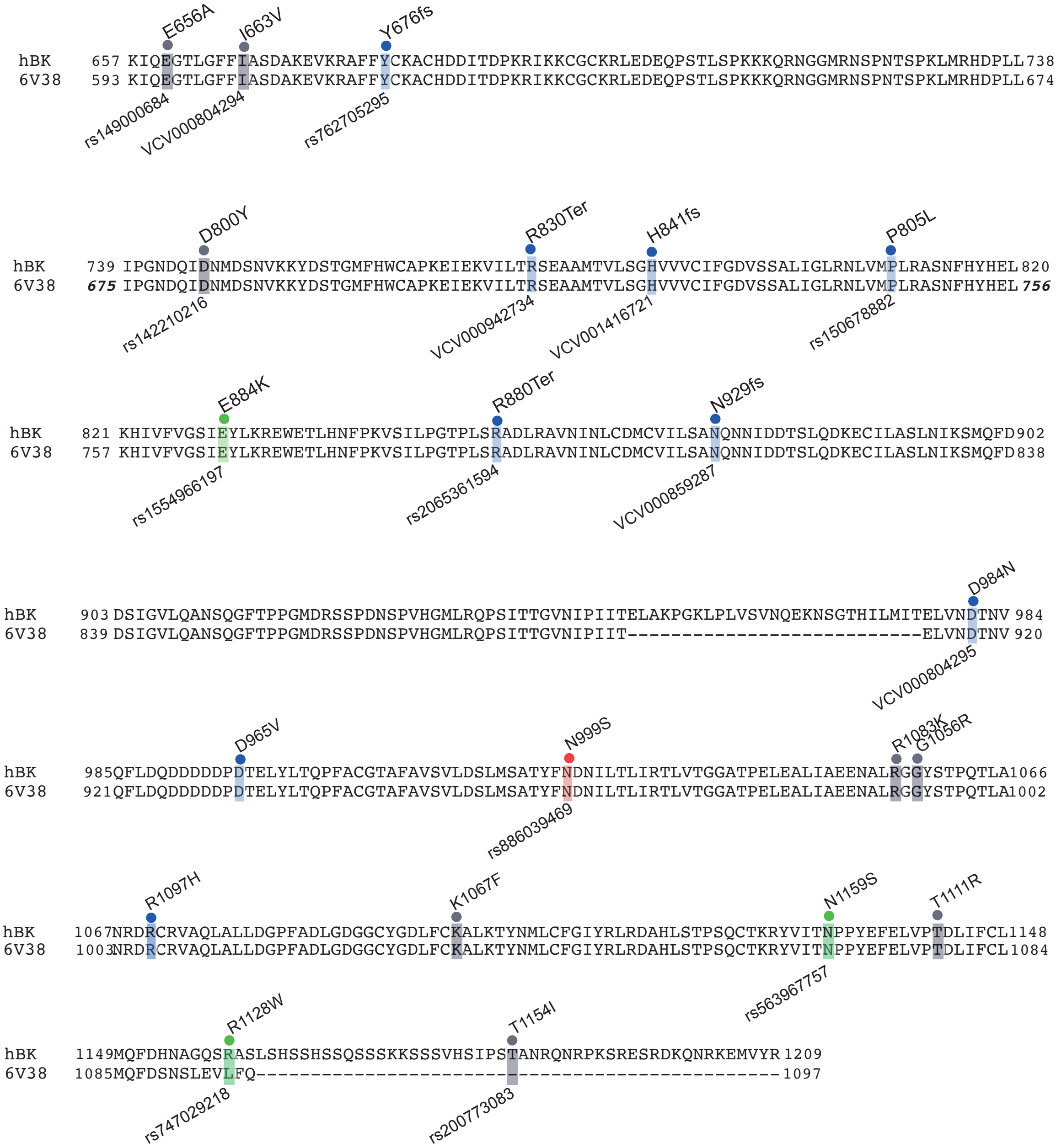

### Supplemental Figure 2

##### Supplemental Figure 2

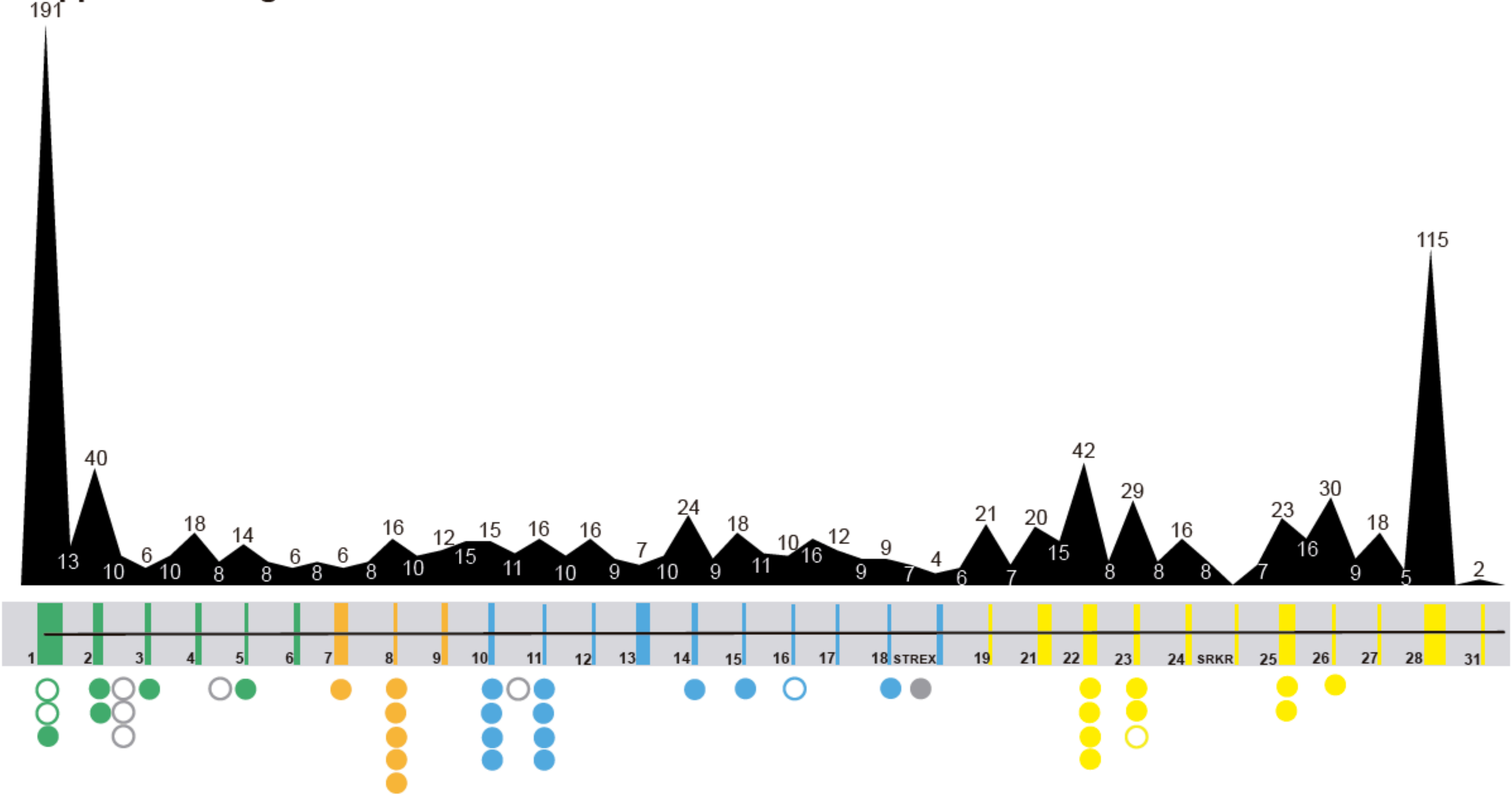
