## Supplemental Table 1 for "Structural mapping of patient-associated *KCNMA1* gene variants"

A

| | SIFT | Polyphen | CADD/PHRED | MetaLR | M-CAP | Mutpred | REVEL |
|---|---|---|---|---|---|---|---|
| **FN** | 10 | 0 | 7 | 9 | 0 | 4 | 1 |
| **FP** | 3 | 4 | 2 | 0 | 4 | 2 | 1 |
| **TN** | 1 | 0 | 2 | 4 | 0 | 2 | 3 |
| **TP** | 6 | 16 | 11 | 8 | 16 | 12 | 17 |

B

| | CADD/Phred | MetaLR | Mutpred | REVEL | WSM | KMS |
|---|---|---|---|---|---|---|
| **A172T** | 2.79 | 0.34 | 0.70 | 0.90 | 4.32 | 4.48 |
| **S351Y** | 2.74 | 0.96 | 0.82 | 0.95 | 5.62 | 5.78 |
| **G354S** | 2.92 | 0.63 | 0.76 | 0.94 | 5.01 | 5.17 |
| **G356S** | 2.92 | 0.63 | 0.76 | 0.94 | 5.01 | 5.17 |
| **G375R** | 2.99 | 0.44 | 0.79 | 0.97 | 4.83 | 5.36 |
| **C413Y** | 3.00 | 0.55 | 0.59 | 0.72 | 4.05 | 4.83 |
| **D434G** | 2.13 | 0.13 | 0.59 | 0.63 | 3.02 | 3.18 |
| **H444Q** | 2.26 | 0.28 | 0.61 | 0.55 | 3.08 | 3.47 |
| **K518N** | 2.34 | 0.09 | 0.47 | 0.14 | 1.44 | 1.60 |
| **N536H** | 2.74 | 0.37 | 0.64 | 0.67 | 3.63 | 5.13 |
| **G567S** | 2.97 | 0.42 | 0.71 | 0.71 | 3.75 | 4.46 |
| **I663V** | 2.46 | 0.79 | 0.58 | 0.67 | 4.22 | 4.39 |
| **P805L** | 2.73 | 0.39 | 0.00 | 0.62 | 2.69 | 2.85 |
| **E884K** | 3.20 | 0.43 | 0.41 | 0.56 | 3.18 | 3.71 |
| **D984N** | 2.94 | 0.41 | 0.38 | 0.37 | 2.55 | 3.08 |
| **N999S** | 2.60 | 0.81 | 0.36 | 0.71 | 4.07 | 5.57 |
| **R1083K** | 2.30 | 0.70 | 0.36 | 0.57 | 3.23 | 3.59 |
| **R1097H** | 2.65 | 0.80 | 0.37 | 0.71 | 4.07 | 4.24 |
| **R1128W** | 2.94 | 0.61 | 0.00 | 0.50 | 2.72 | 2.88 |
| **N1159S** | 2.23 | 0.15 | 0.14 | 0.40 | 1.80 | 1.96 |
