## Supplemental Table 2 for "Structural mapping of patient-associated *KCNMA1* gene variants"

| HP variants | NR | NI | AE (KJ/mol) |
|---|---|---|---|
| S11G17Del | ND | ND | ND |
| G20D | ND | ND | ND |
| I29V | ND | ND | ND |
| L102fs | ND | ND | ND |
| E155Q | ND | ND | ND |
| A172T | 8 | 2 | 12 |
| T176S | 5 | 2 | 34 |
| L226F | 8 | 2 | 34 |
| N237K | 3 | 0 | 0 |
| L271F | 9 | 1 | 23 |
| A314T | 11 | 4 | 46 |
| S351Y | 6 | 2 | 12 |
| G354S | 3 | 0 | 0 |
| G356S | 3 | 1 | 6 |
| G375R | 7 | 2 | 34 |
| C413Y | 11 | 4 | 46 |
| L429fs | ND | ND | ND |
| D434G | 5 | 0 | 0 |
| H444Q | 11 | 1 | 23 |
| N449fs | ND | ND | ND |
| K457E | 5 | 2 | 26 |
| R458Ter | ND | ND | ND |
| I512V | 15 | 3 | 40 |
| K518N | 4 | 0 | 0 |
| A532V | 7 | 2 | 40 |
| N536H | 4 | 2 | 12 |
| C550F | 11 | 3 | 40 |
| G567S | 11 | 1 | 17 |
| M578T | 9 | 1 | 6 |
| E656A | 4 | 0 | 0 |
| I663V | 11 | 0 | 0 |
| Y676fs | ND | ND | ND |
| E736K | ND | ND | ND |
| D800Y | ND | ND | ND |
| P805L | 14 | 1 | 6 |
| R830Ter | ND | ND | ND |
| H841fs | ND | ND | ND |
| R880ter | ND | ND | ND |
| E884K | 6 | 2 | 34 |
| N929fs | 10.5 | 5.5 | 50 |
| D965V | 2 | 0 | 0 |
| D984N | 8 | 1 | 17 |
| N999S | 11 | 1 | 6 |
| G1056R | 6 | 0 | 0 |
| K1067F | 12 | 4 | 71 |
| R1083K | 10 | 0 | 0 |
| R1097H | 16 | 0 | 0 |
| T1111R | 6 | 0 | 0 |
| R1128W | ND | ND | ND |
| T1154I | ND | ND | ND |
| N1159S | ND | ND | ND |
