## Supplemental Figure Legends for "Structural mapping of patient-associated *KCNMA1* gene variants"

### Supplemental Legends

**Supplemental Figure 1. Reference sequence alignment and HP variant database identifiers for 53 variants localized on the BK channel structure.** hBK: NM_001161352.2 and cryo-EM (PDB file 6V38) (Tao and MacKinnon, 2019b). Reference sequence identifier (rs#) or Clinvar record (VCV#) for GOF (red), LOF (blue), and VUS (gray).

**Supplemental Figure 2. KCNMA1 gene variants.** Gene-level depiction for 1068 (synonymous and nonsynonymous) *KCNMA1* variants. 791 exonic variants, with the number in each individual exon in black, and 277 intronic variants (white) annotated above the gene schematic. Of these, 41 variants are either patient-associated or predicted pathogenic in ClinVar (filled and open circles, respectively), or variants of uncertain significance (47 variants, not depicted). Exon coloring scheme: green (voltage sensing domains), orange (pore), blue (RCK1), and yellow (RCK2).

**Supplemental Table 1. Parameters for individual pathogenicity algorithms.** (A) False negative (FN), false positive (FP), true positive (TP), and true negative (TN) parameters for algorithm weighted score (WSM). The FN, FP, and TP for each algorithm are calculated with the cutoffs from the training data set. (B) Individual algorithm scores for the training data set of pathogenic and benign variants (gray). Significant scores are higher than the experimentally-determined cutoffs (CADD/Phred, 2.7; MetaLR, 0.71; Mutpred, 0.43; REVEL, 0.51; and WSM, 2.6).

**Supplemental Table 2. Structural components for the KMS algorithm.** Structural components calculated as the average from the cryoEM structures obtained in the presence and absence of calcium (PDB ID: 6V3G and 6V38). NR, number of residues; NI, number of interactions; AE, average interaction energy; ND, residues not present in the structure.
